# Resolving mechanisms of immune-mediated disease in primary CD4 T cells

**DOI:** 10.1101/2020.01.16.908988

**Authors:** C Bourges, AF Groff, OS Burren, C Gerhardinger, K Mattioli, A Hutchinson, T Hu, T Anand, MW Epping, C Wallace, KGC Smith, JL Rinn, JC Lee

## Abstract

Deriving mechanisms of immune-mediated disease from GWAS data remains a formidable challenge, with attempts to identify causal variants being frequently hampered by linkage disequilibrium. To determine whether causal variants could be identified via their functional effects, we adapted a massively-parallel reporter assay for use in primary CD4 T-cells, key effectors of many immune-mediated diseases. Using the results to guide further study, we provide a generalisable framework for resolving disease mechanisms from non-coding associations – illustrated by a locus linked to 6 immune-mediated diseases, where the lead functional variant causally disrupts a super-enhancer within an NF-κB-driven regulatory circuit, triggering unrestrained T-cell activation.

## INTRODUCTION

Hundreds of genetic loci have been implicated in autoimmune and inflammatory diseases, but the mechanisms by which these effect disease remain largely unknown^1^. An important first step in uncovering these mechanisms is to reduce associated haplotypes down to specific causal variants, whose biological effects mediate disease risk, but statistical attempts to do this have been frustrated by strong linkage disequilibrium (LD), resulting in only a minority of loci being resolved^2–4^. Other methods have sought to re-weight candidate SNPs using their enrichment within functional genomic elements (e.g. tissue-specific regulatory marks)^5–7^, but these methods do not assess whether SNPs have functional consequences, nor identify the biological effect that contributes to disease. This leaves the majority of GWAS loci either unresolved or unresolvable, and the ambition of identifying disease mechanisms largely unrealised^8^. To compound this challenge, the specific gene(s) that are affected by disease-associated variants have not been confirmed for most loci^1^. Many associated haplotypes, for example, contain multiple genes with little or no evidence for any one being causally involved, while other associations are entirely located within intergenic regions (or “gene deserts”) and are often reported to lack candidate genes. Most GWAS associations are attributable to variation in regulatory rather than coding sequence, with significant enrichment in enhancers, and particularly super-enhancers – large enhancer clusters that are usually cell-type specific and control expression of key genes involved in cell state^9^. Testing individual candidate SNPs for effects on transcription – as a means of refining disease-associated haplotypes to specific functional variants – would bypass the limitations of LD and directly assay the process that mediates disease risk but has previously been laborious and expensive. The development of high throughput assays of enhancer activity, such as massively-parallel reporter assays (MPRAs), has now made this possible. MPRAs simultaneously test the regulatory activity of large numbers of short sequences by coupling each to a barcoded reporter gene^10^. By normalising the RNA barcode counts from transfected cells to their equivalent counts in the input plasmid library, MPRAs have identified genetic variants that modulate expression in various settings^11, 12^. A key feature of MPRAs, however, is that the results are determined by the repertoire of transcription factors within the transfected cells, and so could be misleading if an inappropriate cell-type were used. To date, almost all MPRA studies have been performed in cell lines, in part because these are easy to culture and transfect. It is widely recognised, however, that cell lines are poor surrogates for the types of the primary immune cells that drive autoimmune disease^13–15^.

Here, we adapt an MPRA for use in resting and stimulated primary CD4 T cells – the cell-type whose regulatory DNA is most highly enriched for immune-mediated disease SNPs^2, 16–18^. We use this method to simultaneously test individual candidate SNPs from 14 immune disease-associated gene deserts for expression-modulating activity. By treating the results of this assay as a basis to explore the underlying biology, we gain previously unappreciated insights into the effects of disease-associated variation, as illustrated by the pleiotropic 6q23 locus. At this multi-disease-associated haplotype, we uncover a molecular mechanism whereby a common variant – identified via its expression-modulating effect in CD4 T cells – disrupts an NF-κB-driven pathway that normally limits T cell activation through the dynamic formation of a *TNFAIP3* super-enhancer. Disruption of this feedback circuit releases activated CD4 T cells from an intrinsic molecular brake and thus reveals a mechanism by which a disease-associated haplotype can causally change biology, and a pathway that would appear to be pervasively involved in human autoimmunity.

## RESULTS

### Adaptation of MPRA for use in primary CD4 T cells

To determine whether causal variants could be identified via their functional effects, we designed an MPRA to assess candidate SNPs (all variants with r^2^≥0.8 with the lead SNP) at 14 gene deserts linked to one or more of 10 different immune-mediated diseases (**Fig. 1a**, **Table 1**)^19–27^. Several of these loci cannot be resolved by fine-mapping^2–4^. Gene deserts were selected because: (1) less is known about how these predispose to disease compared to regions containing candidate genes, (2) other non-coding mechanisms (such as splicing effects^28^) are unlikely to account for these associations, and (3) many of these contain epigenetic marks consistent with enhancer activity^9, 18, 29^. To maximise the genomic context tested around each SNP, we designed 3 overlapping constructs for every SNP allele^12^, and synthesised additional oligonucleotides to test combinations of risk alleles if more than one SNP could be assayed within the same construct. We also included oligonucleotides that tiled each locus at 50bp intervals to test for enhancer activity – and enable us to exclude regions that lacked this.

**Figure 1.**
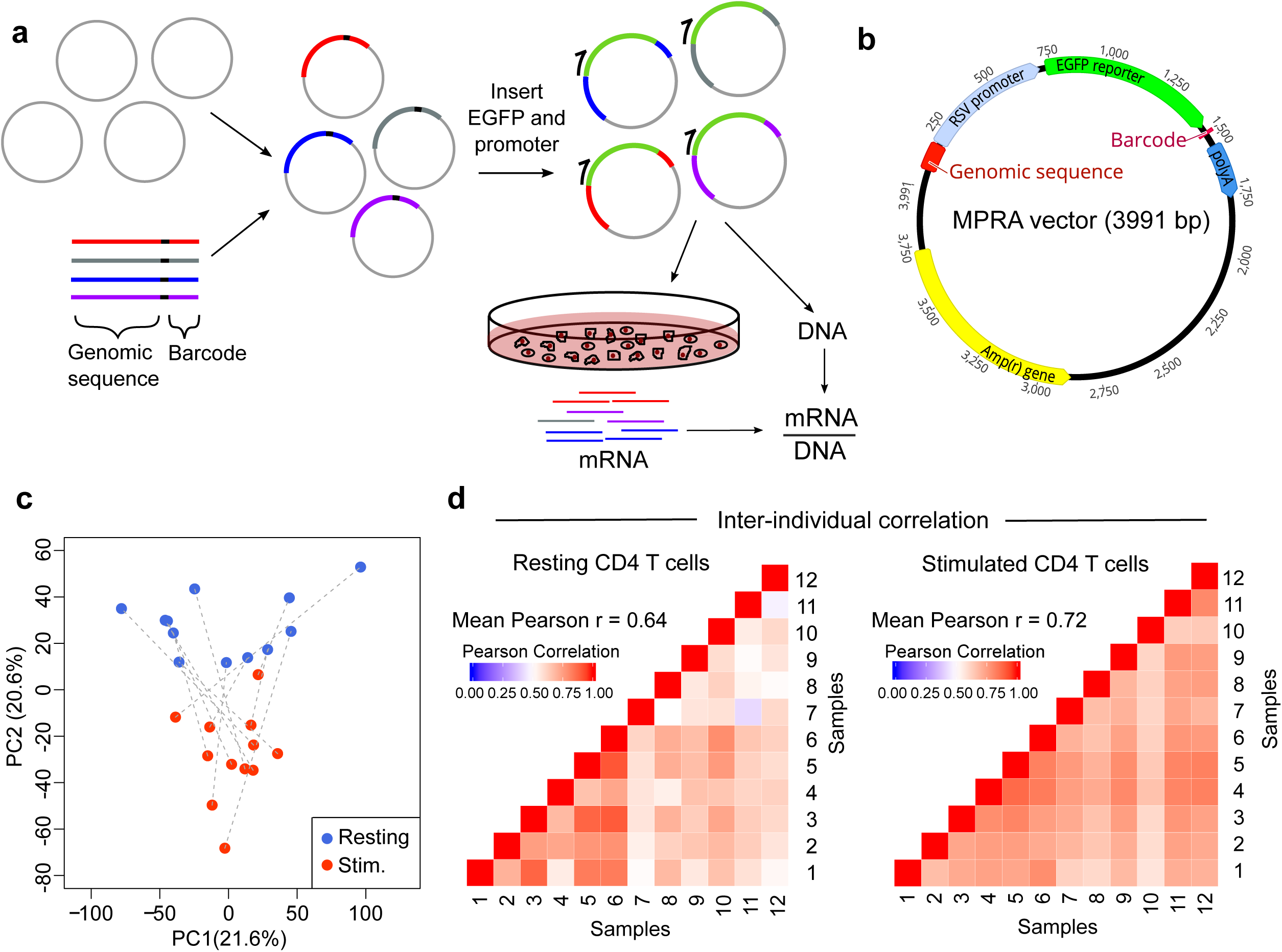
Development of MPRA for use in primary human CD4 T cells. **a** Experimental workflow for MPRA: oligonucleotide library is cloned into an empty vector and a reporter gene and promoter are subsequently inserted using restriction sites within the oligonucleotide. The assembled plasmid is transfected into primary CD4 T cells and RNA is extracted after 24 hours. RNA barcode counts are normalised to their respective counts in the input plasmid library (DNA), which is sequenced separately. **b** Adapted MPRA plasmid incorporating RSV promoter. **c** Principal component analysis of scaled element counts (sum of barcodes tagging same genomic construct in mRNA) in resting and stimulated CD4 T cells from 12 donors. Dotted lines indicate samples from the same donor. **d** Heatmaps showing pairwise comparison of MPRA activity for all constructs (mRNA/DNA) between donors – left panel: resting CD4 T cells; right panel: stimulated CD4 T cells.

**Table 1.** Autoimmune disease associations at 14 gene deserts.

| tag SNP | $r^2$<br>between<br>tag SNPs | Chr. | Associated disease(s) | Distance<br>to nearest<br>gene (kb) | Number<br>of SNPs<br>($r^2 \geq 0.8$) | Haplotype<br>block size<br>(kb) |
| --- | --- | --- | --- | --- | --- | --- |
| rs883220 | - | 1p34 | RA | 102.3 | 9 | 30.2 |
| rs6759298 (AS)<br>rs10865331 (Ps) | 0.86 | 2p15 | AS, Ps | 99.5 | 12 | 33.9 |
| rs1534422 | - | 2p24 | ATD | 208.2 | 12 | 15.9 |
| rs1813375 | - | 3p24 | MS | 203.9 | 15 | 10.9 |
| rs2611215 | - | 4q32 | T1D | 139.4 | 20 | 16.7 |
| rs12186979 | - | 5p13 | AS | 59.7 | 5 | 97.8 |
| rs17119 | - | 6p23 | UC, CD, MS | 512.1 | 44 | 22.6 |
| rs2327832 (SLE)<br>rs6920220 (rest)<sup>a</sup> | 0.92 | 6q23 | RA, CeD, UC, CD, SLE<sup>b</sup>, T1D<sup>c</sup> | 143.6 | 9 | 47.5 |
| rs1991866 | - | 8q24 | UC, CD | 136.2 | 28 | 22.0 |
| rs2456449 | - | 8q24 | MS | 219.3 | 17 | 19.5 |
| rs4409785 | - | 11q21 | ATD, vitiligo<sup>d</sup> | 181.2 | 3 | 9.7 |
| rs1456988 | - | 14q32 | T1D | 1086.9 | 38 | 14.0 |
| rs1297258 | - | 21q21 | UC, CD | 274.0 | 38 | 24.1 |
| rs2836883 (AS, PSC)<br>rs2836878 (UC, CD) | 1.0 | 21q22 | AS, PSC, UC, CD | 87.1 | 13 | 5.8 |

Genetic associations were identified from published immunochip data. For each locus, the extended haplotype (LD region tagged by all SNPs with $r^2 \geq 0.8$ and extended by 50kb on either side) does not contain coding or well-characterised non-coding genes. Haplotype block size represents region tagged by all SNPs with $r^2 \geq 0.8$ .
<sup>a</sup> CeD tag SNP rs17264332 ( $r^2 = 1.0$ with rs6920220).
<sup>b</sup> Association reported subsequently<sup>64</sup>.
<sup>c</sup> Association reported using $P < 1 \times 10^{-5}$ to obtain a Bayesian posterior probability for T1D association given known associations with other diseases<sup>22</sup>.
<sup>d</sup> Vitiligo association reported in GWAS, not immunochip<sup>65</sup>.
SNP, Single Nucleotide Polymorphism; Chr., chromosome; AS, Ankylosing Spondylitis; Ps, Psoriasis; PSC, Primary Sclerosing Cholangitis; UC, ulcerative colitis; CD, Crohn's disease; RA, rheumatoid arthritis; CeD, coeliac disease; SLE, Systemic Lupus Erythematosus; T1D, Type 1 Diabetes; ATD, autoimmune thyroid disease; MS, multiple sclerosis.

After assembly, the MPRA plasmid library was transfected into primary CD4 T cells from healthy donors (**Fig. 1a, Methods**) but no expression of the reporter gene was detected (**Figs. S1a, S1b**). After confirming that successful transfection had occurred (**Fig. S1b**), we surmised that the minimal promoter, which is conventionally used for MPRA, may be insufficient to initiate transcription in primary T cells. In cell line-based MPRA studies, stronger promoters have been shown to produce highly comparable results to those obtained using a minimal promoter^30, 31^. We therefore screened a series of promoters in CD4 T cells (**Fig. S1c**) and selected the Rous Sarcoma Virus (RSV) promoter for incorporation into an adapted MPRA vector (**Fig. 1b**) as this robustly initiated transcription but was not so strong as to preclude further amplification.

After assembly, the adapted MPRA plasmid library was transfected into resting and stimulated CD4 T cells from 12 healthy donors. Multiple biological replicates (donors) were used to ensure that the results were reproducible, and control for inter-individual differences in CD4 T cell composition and the reduced dynamic range expected with a stronger promoter. After 24 hours, GFP was detected and RNA was extracted to quantify expression of each barcode using high-throughput sequencing (**Fig. S1d**). Following pre-processing, the barcode counts were collapsed to individual genomic constructs for further analysis (**Methods**). Using principal component analysis, we found that the activation state of T cells was responsible for much of the total variance (**Fig. 1c**) and that the transcriptional activity of constructs – obtained by normalising the RNA barcode counts to their respective counts in plasmid library – correlated well between different individuals (**Fig. 1d**). To detect expression-modulating variants, we used QuASAR-MPRA^32^ (**Fig. 2a**) and combined the results from each donor using a fixed-effects meta-analysis (**Methods**). Significant expression-modulating activity was detected at one or more constructs for 8/10 positive control SNPs (comprising 5 known expression-modulating variants^11^, 2 single variant eQTLs^2^, and 3 synthetic SNPs that included / disrupted a binding site for a transcription factor active in T cells) (**Fig. 2b**). Enhancer activity was also detected in the positive control regions for the tiling analysis, while no such activity was detected in the negative controls (**Table S1, Methods**). To validate the observed effects, we tested the most significant expression-modulating SNP at each haplotype, 2 positive control SNPs and 5 SNPs with no allele-specific effects using a complementary luciferase-based system (**Fig. 2c**). Despite using a different promoter and quantification method, we observed a strong correlation between the MPRA and luciferase results (Pearson r = 0.87) – indicative of genuine expression-modulating effects that are likely to be physiologically relevant (**Fig. 2d**). Altogether, these data indicate that MPRA can be adapted for use in primary CD4 T cells, and that the results reflect the activation state of the cells and can identify constructs with known regulatory effects.

**Figure 2.**
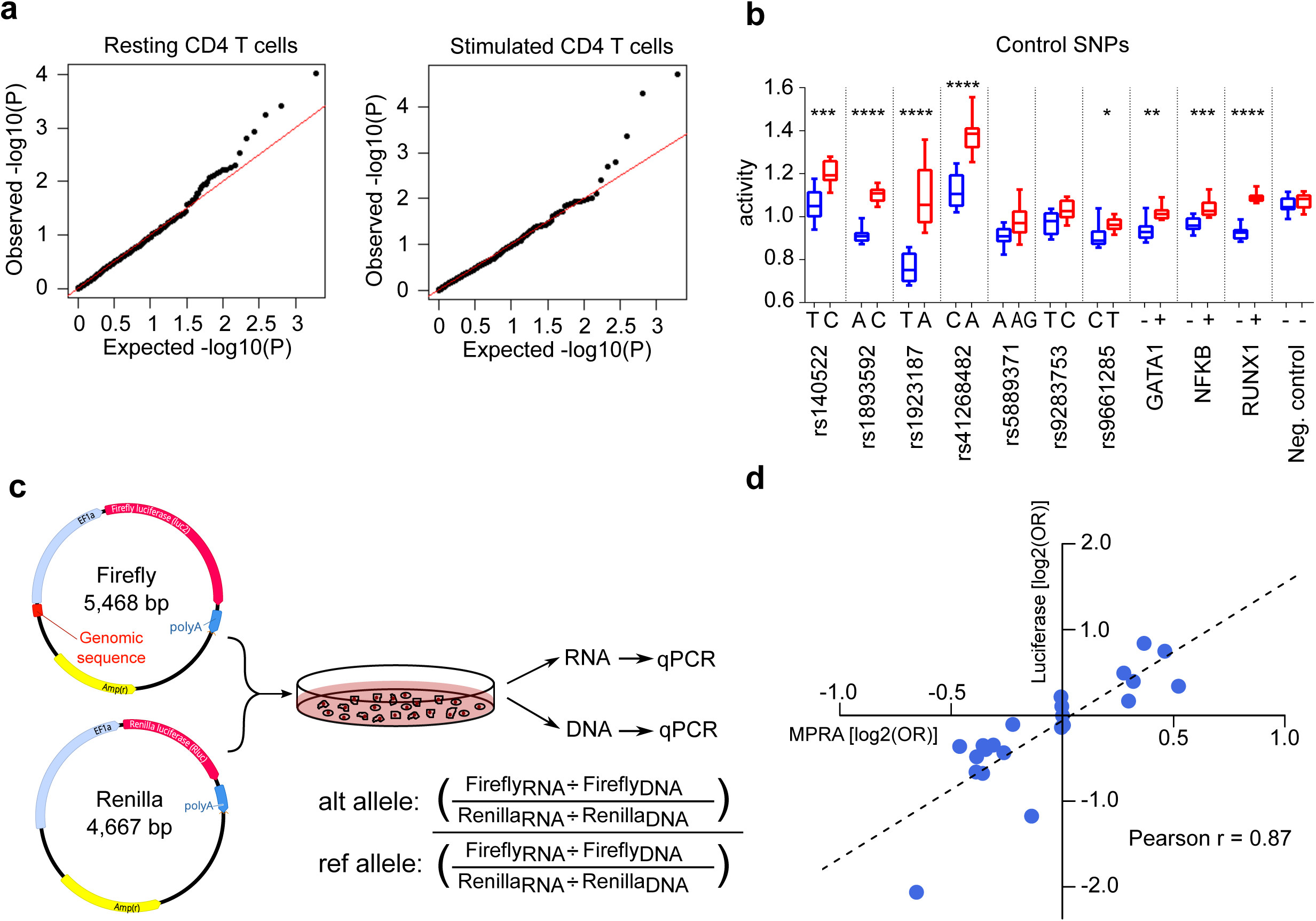
Allele-specific expression-modulating effects in CD4 T cells. **a** qq plots of the observed -log10(P) values versus the expected -log10(P) values under the null hypothesis for representative resting and stimulated CD4 T cell samples. **b** Activity of each allele at 10 positive control SNPs and 1 negative control SNP in stimulated T cells. GATA1, NF-κB, and RUNX1 constructs were designed to include a binding site for the indicated transcription factors (+) or with that site disrupted (-). FDR-corrected statistical significance is shown (fixed effects meta-analysis P value): *<0.05; **<0.01, ***<0.001, ****<0.0001. Box and whisker plots represent median and IQR (box) and min-to-max (whiskers). **c** Experimental workflow for validation experiment using a different promoter (EF1α), reporter gene (luciferase) and quantification method (qPCR). **d** Expression-modulating effect of each SNP [log2(OR)] as measured in MPRA and validation experiments. OR were calculated using the median activity of allelic constructs and are presented with respect to the risk allele.

### Adapted MPRA in CD4 T cells provides insights into the biological effects of genetic associations

After establishing that MPRA could be performed in primary CD4 T cells, we next examined the results at disease-associated loci (**Figs. S2, S3, Tables S2, S3**). To determine whether adapted MPRA would identify known causal variants, we used an inflammatory bowel disease (IBD)-associated locus that has previously been fine-mapped to a single variant, rs1736137 (ref 3). In both resting and stimulated T cells, this SNP had highly significant expression-modulating activity, with the IBD-risk allele consistently increasing transcription (**Fig. 3a**). As a further proof of principle, we next examined an ankylosing spondylitis-associated locus, where Bayesian fine-mapping using corrected coverage estimates resolves the association to 3 SNPs in the 99% credible set (rs6759298, rs4672505 and rs13001372). In stimulated T cells, one of these SNPs (rs6759298) had the most significant expression-modulating effect of all the SNPs tested at this locus, while the others had negligible effects on transcription (**Fig. 3b**) – thus identifying the variant that could causally change biology.

**Figure 3.**
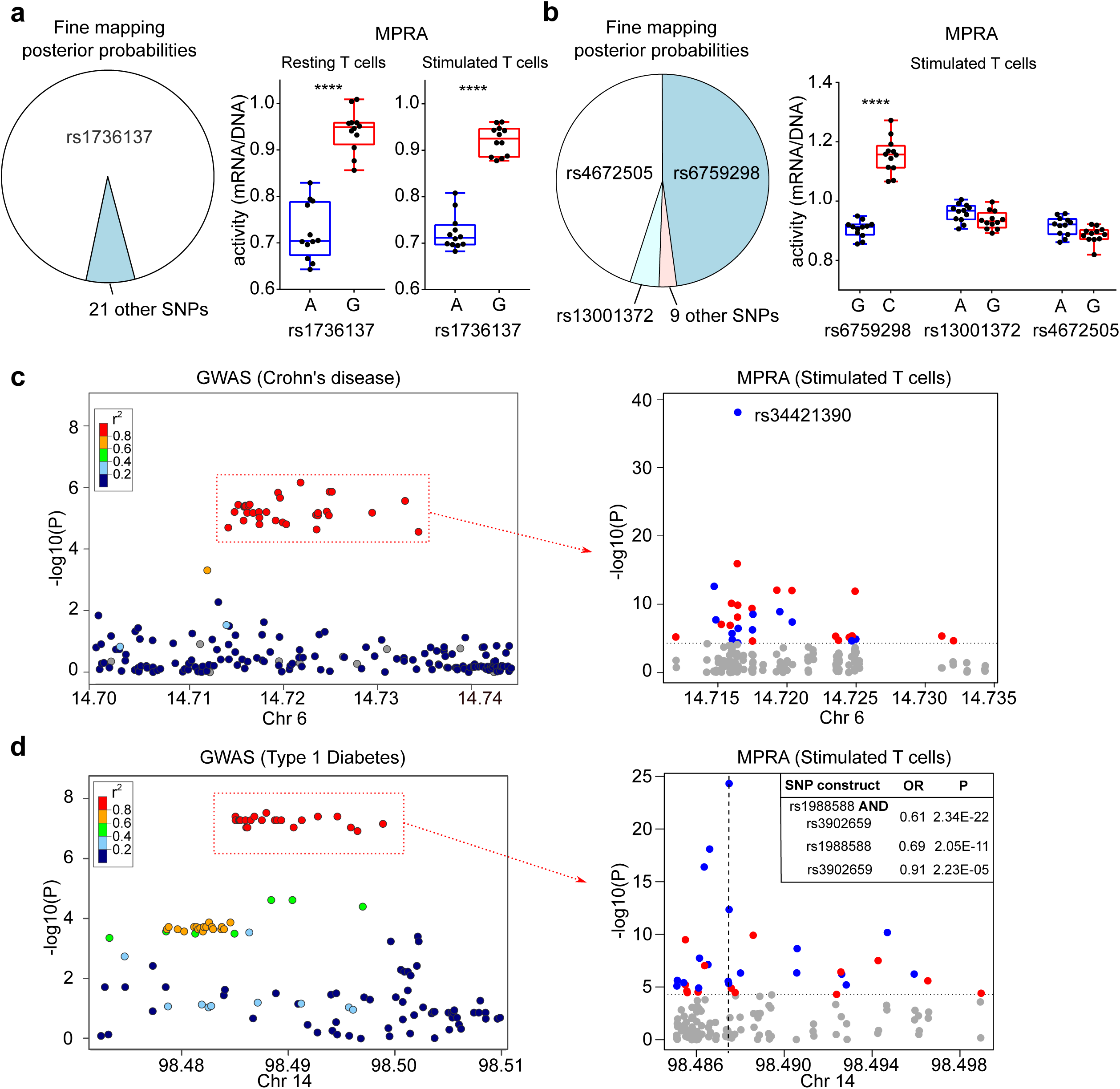
MPRA in CD4 T cells identifies biological effects of disease associations. **a** Pie chart depicting fine-mapping results^3^ (posterior probabilities) for an IBD-associated locus on 21q21, with rs1736137 assigned an 88% posterior probability of being causal (left panel). MPRA results in resting (centre panel) and stimulated CD4 T cells (right panel) showing that rs1736137 has a significant expression-modulating effect. **b** Pie chart depicting Bayesian fine mapping results for an AS-associated locus on 2p15 (left panel). MPRA results in stimulated T cells (right panel) showing that rs6759298 has a significant expression-modulating effect (the strongest of any variant at this locus) while the other candidate SNPs have negligible effects. **c** GWAS results at a Crohn’s disease and multiple sclerosis-associated locus on 6p23 (data from ref 62; left panel). MPRA for candidate SNPs in stimulated T cells (right panel) identifies a single SNP (rs34421390) with by far the greatest expression-modulating effect at this locus, where the risk allele reduces expression (blue, risk allele reduces expression; red, risk allele increases expression). Dotted horizontal line represents significance threshold (corrected for multiple-testing). **d** GWAS results at a Type 1 Diabetes-associated locus on 14q32 (data from ref 22; left panel). MPRA for the candidate SNPs in stimulated T cells (right panel) identifying that the construct with the largest and most significant effect contains the risk alleles for 2 SNPs (rs1988588 and rs3902659), each of which has a smaller concordant effect when tested individually (position indicated by vertical dotted line). Box and whisker plots represent median and interquartile range (box) and min to max (whiskers). **** FDR-corrected meta-analysis P<0.0001.

We next investigated whether adapted MPRA could resolve possible causal variants at other loci, and so provide testable hypotheses into disease mechanisms. Specific variants with strong functional effects were identified at several loci (**Figs S2, S3**), including – for example – a chromosome 6 locus that is associated with both IBD and multiple sclerosis. Of 44 candidate SNPs within the shared disease-associated haplotype, a single variant (rs34421390) had by far the largest and most significant expression-modulating effect in both resting and stimulated CD4 T cells (**Fig. 3c**). This provides a focus for studying the upstream biology, and also demonstrates that the risk haplotype reduces transcription – an important finding since the locus interacts with the promoter of *JARID2*, a component of the Polycomb-Repressive Complex 2, in CD4 T cells^33^.

We made similar insights at a Type 1 Diabetes-associated locus, which contains 38 SNPs in strong LD. At this haplotype, the largest and most significant expression-modulating effect occurred with a construct containing the risk alleles for two adjacent SNPs (rs1988588 and rs3902659) which are located 60bp apart (**Fig. 3d**). Both SNPs had similar effects when tested individually (with the risk allele reducing transcription) but these were weaker than with the construct containing both risk alleles (**Fig. 3d**). This raises the possibility that the functional effect of this haplotype is mediated by a synergistic interaction between two adjacent SNPs, rather than a single causal variant – a prospect that could not be derived from genetic data since the SNPs are in complete LD (r^2^>0.9999, ref 22). This provides a basis to study the molecular mechanisms at this locus, which could help resolve the underlying biology.

Altogether, these results demonstrate that MPRA in primary CD4 T cells can identify variants that causally alter gene expression, and so provide testable hypotheses into possible disease mechanisms, while simultaneously identifying the nature of the functional effect.

### MPRA identifies an expression-modulating variant that disrupts NF-κB binding and super-enhancer formation

To confirm that MPRA in primary CD4 T cells could help resolve disease mechanisms, we selected a pleiotropic locus on chromosome 6 for further study (**Fig. 4a**). This region was chosen for several reasons. First, it was the only haplotype that was associated with 6 different diseases (**Table 1**), highlighting the biological importance of the locus^19, 20, 22, 26, 34^. Second, despite receiving considerable attention, there is still uncertainty regarding the causal gene at this locus, with some studies implicating *TNFAIP3*, mainly because this is the closest plausible candidate^19, 22, 35, 36^ while others suggest that *IL20RA* is responsible^37, 38^. Third, statistical fine-mapping has been attempted at this locus but has been hampered by strong LD^2, 3^ (**Fig. 4b**). In the MPRA, the same SNP (rs6927172) showed the strongest expression-modulating effect in both resting and stimulated T cells, with the risk allele consistently reducing transcription (**Figs. 4c, S2, S3**). Further examination of this SNP revealed that it lies in a highly conserved region (**Fig. S4a**) containing an experimentally-validated NF-κB binding motif, to which all NF-κB dimers can bind. rs6927172 is located at position 10 within this common 11-mer binding site, with the risk allele predicted to disrupt binding (**Fig. 4d**). Consistent with this, *in silico* methods, including Deepsea^40^, also predicted that this SNP would disrupt NF-κB binding (**Fig. S4b**). To determine whether allele-specific NF-κB binding might account for the MPRA result, we transfected the MPRA plasmid library into CD4 T cells, immunoprecipitated NF-κB, and quantified the plasmids to which it was bound. We confirmed that NF-κB differentially bound to rs6927172-containing plasmids in a manner consistent with the MPRA result and *in silico* prediction (**Fig. S4c**). We therefore investigated whether allele-specific NF-κB binding might also occur at the native locus in primary CD4 T cells. To do this, we isolated CD4 T cells from healthy donors who were heterozygous at rs6927172 and immunoprecipitated NF-κB to quantify the relative binding to each allele using an adapted genotyping assay (**Methods**). We observed that in stimulated T cells, NF-κB exhibited allele-specific binding with reduced binding to the rs6927172 risk allele – consistent with the MPRA result (**Fig. 4e**). Conversely, we did not detect allele-specific binding in resting T cells, which may reflect insufficient NF-κB signalling and suggests that the MPRA result in resting cells could be partly due to the transient activation that can occur following nucleofection^41^.

**Figure 4.**
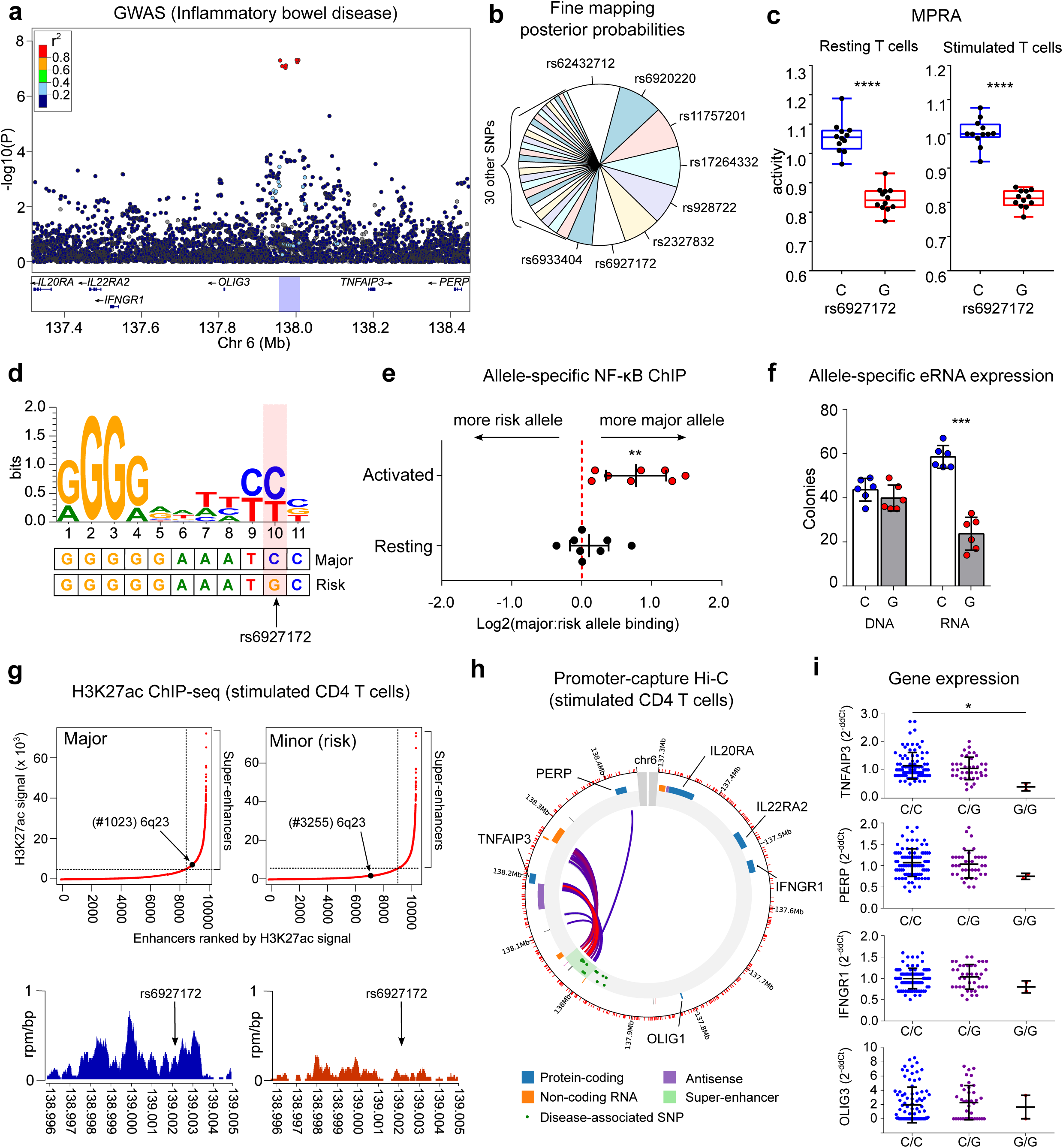
MPRA in CD4 T cells identifies an expression-modulating variant that disrupts NF-κB binding and enhancer function. **a** IBD GWAS results^62^ at a multi-disease-associated locus on chromosome 6q23. **b** Fine-mapping results^3^ (posterior probabilities) for candidate SNPs at this locus. **c** A single variant (rs6927172) has the largest and most significant expression-modulating activity in resting (left panel) and stimulated CD4 T cells (right panel) with the risk allele reducing transcription. Plots represent median and IQR (box) and min-to-max (whiskers). FDR-corrected meta-analysis P value shown. **d** Sequence logo for an experimentally-validated NF-κB binding motif. The genomic sequence around rs6927172 is aligned below. **e** Allele-specific NF-κB binding in CD4 T cells from rs6927172 heterozygotes, demonstrating reduced NF-κB binding to the risk allele following stimulation (n=8; one-sample *t*-test, two-tailed). **f** Allele-specific expression of enhancer RNA in heterozygous CD4 T cells. DNA represents technical control (n=6; paired *t*-test; two-tailed). **g** Genome-wide H3K27ac ChIP-seq in stimulated CD4 T cells from major- and minor-allele homozygotes at rs6927172 (n=6). Upper panels show input-normalised H3K27ac signals (generated by ROSE^46^) plotted against enhancer rank. Super-enhancers are conventionally defined above the inflection point of the curve. Lower panels show H3K27ac reads from a major-(left) and a minor (risk) allele homozygote (right) in a 9kb window around rs6927172. **h** Promoter-capture Hi-C overview plot depicting interactions of the 6q23 super-enhancer. Data from ref 33. **i** Expression of genes on 6q23 in CD4 T cells from 131 patients with active IBD, stratified by rs6927172 genotype (qPCR; one-way ANOVA). Error bars represent SD. *IL20RA* and *IL22RAR2* expression was not detected. Data represent mean+/-SEM, unless indicated. * P<0.05; ** P<0.01, **** P<0.0001.

To determine whether differential NF-κB binding might affect enhancer strength, we exploited the fact that active enhancers are transcribed, producing enhancer-(e)RNAs whose abundance generally correlates with enhancer activity^42, 43^. Using an allele-specific expression assay, we compared the amount of eRNA transcribed from each allele in stimulated heterozygous CD4 T cells – thus ensuring that external factors would affect both alleles equally^44^. We confirmed allele-specific expression of the eRNA at this locus, with significantly less transcription from the risk allele, in which the NF-κB binding site is disrupted (**Fig. 4f**). This suggests that the effect of the disease-associated haplotype is to diminish enhancer activity by perturbing an NF-κB binding site – potentially linking the genetic association to a specific functional deficit.

During inflammatory responses, NF-κB binding has been reported to direct dynamic super-enhancer formation^45^. We therefore sought to better characterise the functional consequences of allele-specific NF-κB binding at this locus. To do this, we performed histone H3K27 acetylation (H3K27ac) ChIP-sequencing in stimulated CD4 T cells from major and minor allele homozygotes at rs6927172. This facilitated a genome-wide comparison of active regulatory regions, and enabled us to better characterise the effect of rs6927172 on enhancer activity. We observed consistently stronger enhancer activity at this locus in major allele homozygotes compared to minor (risk) allele homozygotes (**Fig. S5a**). To improve peak-calling and generate representative datasets, we combined the genotypic replicates for subsequent analysis. Using the Rank Ordering of Super-Enhancers (ROSE) algorithm^46^, we found that rs6927172 was located within a 45.5kb super-enhancer in major allele homozygotes (**Figs. 4g, S5a**). This super-enhancer appears to be T cell-specific, and potentially activation-specific, since it has also been detected in stimulated Th17 cells, but not in 27 other primary tissues nor 5 other immune cell types^9^. Consistent with this, we found that many of the transcription factors predicted to bind within the constituent elements of the super-enhancer were involved in T cell activation (**Figs. S5b, S5c**). In contrast to the strong enhancer activity in major allele homozygotes, there was negligible enhancer activity at the rs6927172 locus in minor allele homozygotes (**Figs. 4g, S5a**). Indeed, while ROSE analysis identified preserved enhancer activity 1.5kb upstream and 18.8kb downstream of this SNP (extending to the 5’ and 3’ ends of the annotated super-enhancer) the overall enhancer strength across this region was four-fold lower in the presence of the risk allele, and super-enhancer formation was accordingly disrupted (**Figs. 4g, S5a**).

To understand why disrupting the formation of an NF-κB-driven super-enhancer might predispose to multiple immune-mediated diseases, we next investigated the genes that it regulated. Using available promoter-capture Hi-C data from stimulated CD4 T cells, we confirmed that the majority of super-enhancer interactions were either with the promoter of *TNFAIP3* or with a region ∼41kb downstream of *TNFAIP3* that also interacts with the *TNFAIP3* promoter (**Fig. 4h**). Consistent with this, we found that *TNFAIP3* expression in primary CD4 T cells from 131 patients with active IBD correlated with rs6927172 genotype, whereas no such correlation was observed for other genes at this locus (**Fig. 4i**). Expression of *IL20RA*, which was suggested to be causal based on experiments in cell lines^37, 38^, could not be detected in primary CD4 T cells, and appears not to be expressed in primary immune cells according to publicly-available datasets^47, 48^ (**Fig. S5d**). In contrast, *TNFAIP3* is highly expressed in effector CD4 T cell lineages^47^ (**Fig. S5e**) and encodes A20, a key negative regulator of NF-κB signalling and an early target gene of NF-κB^49, 50^.

Collectively, these data are consistent with a model in which NF-κB signalling in stimulated CD4 T cells leads to the formation of a super-enhancer at an immune-mediated disease locus that drives expression of a key NF-κB inhibitor – thereby limiting inflammatory responses. This regulatory circuit can be disrupted by a common expression-modulating variant, such that NF-κB binding and enhancer activity are diminished in the presence of the risk allele. This would be predicted to lead to excessive inflammatory responses in CD4 T cells, consistent with an association with multiple immune-mediated diseases.

### The NF-κB binding site disrupted by rs6927172 regulates *TNFAIP3* expression and inflammatory responses in CD4 T cells

To test whether our proposed model was correct, we investigated the consequences of deleting the NF-κB binding site in primary CD4 T cells using CRISPR-Cas9. Efficient genome editing in primary T cells usually requires the cells to be pre-activated^51, 52^, but a method was recently described for editing resting T cells^53^. Since we wished to study the effects of editing upon subsequent T cell activation, we similarly optimised conditions for editing resting CD4 T cells (**Methods**, **Fig. S6a**) – achieving on-target indels in up to 80% of cells (depending on the gRNA) and efficient knock-down of surface proteins (**Fig. S6b**). We therefore designed gRNAs flanking the rs6927172-containing NF-κB binding site and used these in different combinations to reduce the chance that observed effects were due to off-target activity (**Fig. 5a**). We observed mean editing rates of 60-70% for 3 of the 4 combinations of gRNAs, of which ∼80% of predicted indels ablated the NF-κB binding site (**Fig. 5b**). Of note, the lower editing rate observed with the fourth gRNA combination probably reflects steric hindrance between Cas9 molecules since the offset between the gRNAs was only 4bp^54^.

**Figure 5.**
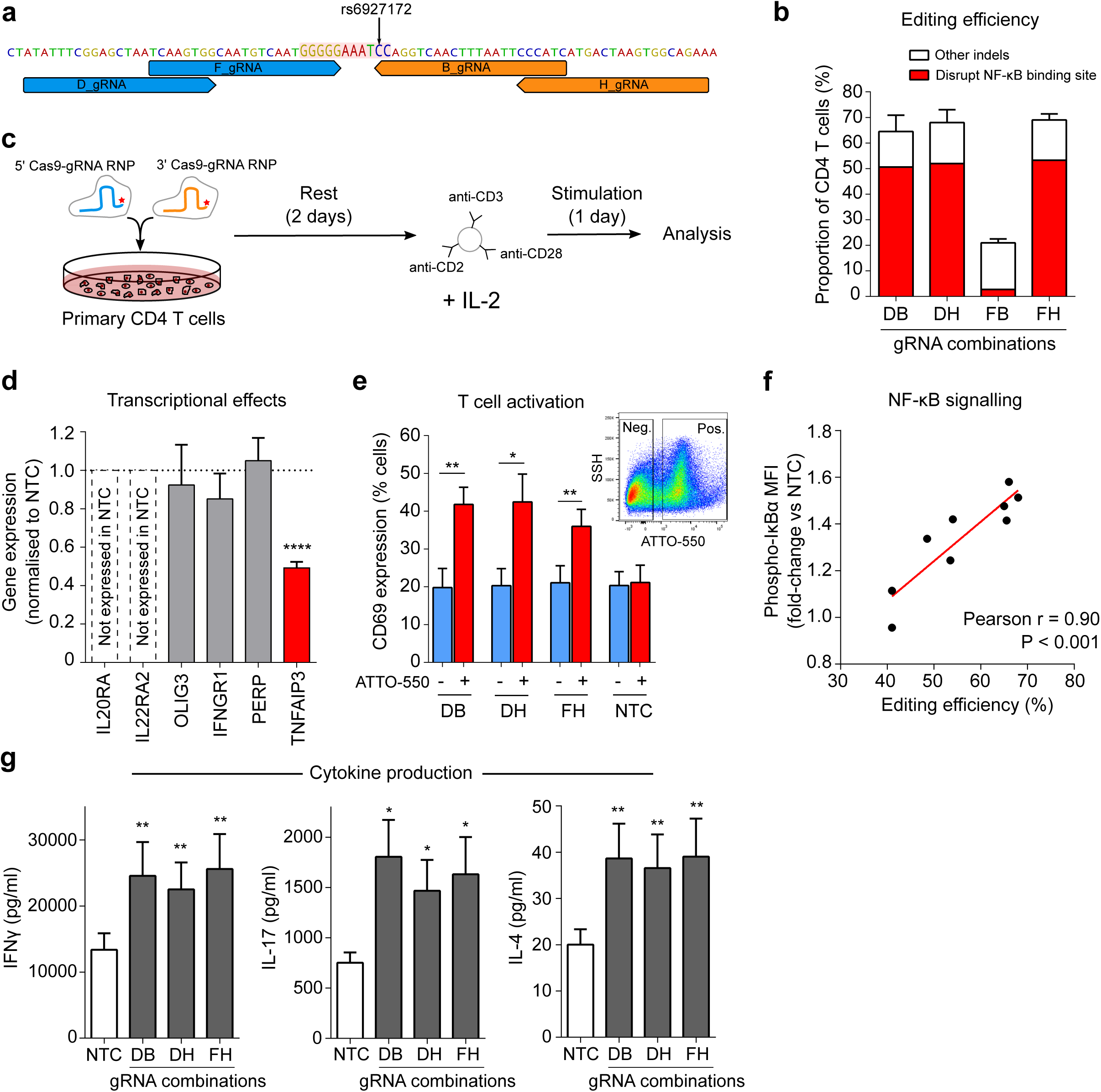
Deletion of the NF-κB binding site, disrupted by rs6927172, dysregulates *TNFAIP3* expression and increases CD4 T cell activation. **a** Location of gRNAs flanking the NF-κB binding site (highlighted). **b** Editing efficiency at the target site in primary CD4 T cells, for indicated combinations of 5’ and 3’ gRNAs (n=6 for DB, DH and FH, and n=2 for FB – stopped due to poor efficiency). Distribution of indels assessed using ICE^63^. **c** Experimental workflow: equimolar amounts of 5’ and 3’ gRNA-containing RNPs (fluorescently tagged with ATTO-550) were nucleofected into unstimulated CD4 T cells, which were rested for 48 hours before stimulation with anti-CD2/3/28 microbeads and IL-2 for 24 hours. **d** Expression of genes on 6q23 in EU-containing mRNA (EU added at time of stimulation) showing that deletion of the NF-κB binding site specifically reduces transcription of *TNFAIP3*, but not other genes at this locus (n=6; one sample t-test). Representative data shown from the DH gRNA combination. **e** Expression of CD69, an activation marker, following CRISPR editing with each gRNA combination or the non-targeting (negative) control (NTC) – data shown for ATTO-550 positive (RNP-containing) and negative cells (n=6; paired t-test, one-tailed). Inset flow cytometry plot depicting representative gating of ATTO-550 positive and negative cells. **f** Correlation between editing efficiency (total indel rate) and levels of phosphorylated IκBα in CD4 T cells (normalised to the mean fluorescence intensity in the NTC). **g** Secretion of effector cytokines following deletion of the NF-κB binding site – reflecting Th1 (left panel, IFNγ), Th17 (centre panel, IL-17A) and Th2 subsets (right panel, IL-4) (n=6, paired t-test, one-tailed). Data represent mean+/-SEM. * P<0.05; ** P<0.01; **** P<0.0001.

We next investigated how deleting the NF-κB binding site would impact transcription locally. After nucleofection with Cas9-gRNA ribonucleoproteins (RNPs), CD4 T cells were rested for 48 hours and then stimulated for 24 hours (**Fig. 5c, Methods**). To specifically quantify RNA that was transcribed during T cell activation – and after editing – we added 5-ethynyl uridine (EU) at the time of stimulation to facilitate nascent RNA capture (**Methods**). RNA that incorporated this modified base was purified and expression of all protein-coding genes within 1.5Mb of the deletion site was measured and normalised to a non-targeting control. Of the six genes tested, only *TNFAIP3* expression was significantly altered (**Fig. 5d**). Moreover, individual deletions of the other candidate SNPs within the disease-associated super-enhancer did not significantly alter *TNFAIP3* expression – consistent with dysregulation of enhancer activity being specific to rs6927172 (**Fig. S6c)**.

To understand the biological consequences of this effect, we next examined markers of T cell activation. Using a fluorescently-tagged gRNA that is detectable by flow cytometry, we distinguished CD4 T cells that contained the Cas9-gRNA RNPs (and were more likely to have been edited) from those that did not. Analysing these populations separately, we observed a specific increase in CD69 expression, an early marker of T cell activation^55^, in the RNP-containing cells, that was not present in the non-targeting control, nor in the RNP-negative cells from the same transfection (**Fig. 5e**). This indicated that deletion of the NF-κB binding site, which is physiologically disrupted by rs6927172, leads to increased T cell activation.

To further explore the underlying mechanism, we used flow cytometry to quantify IκBα phosphorylation, a key step in NF-κB signalling. After normalising to the non-targeting control, the increase in phospho-IκBα was shown to directly correlate with the overall editing efficiency (**Fig. 5f**) – suggesting that NF-κB signalling increases proportionally with deletion of the NF-κB binding site. To understand how this would affect CD4 T cell effector function, we quantified cytokine production and found that deletion of the NF-κB binding site led to increased effector cytokine production from all major T helper cell lineages, consistent with unrestrained inflammatory responses (**Fig. 5g**). Finally, to confirm that these results were consistent with a *TNFAIP3*-dependent effect, we directly disrupted *TNFAIP3* using CRISPR editing and showed that this phenocopied the observed effects, with marked increases in T cell activation (**Fig. S6d**) and effector cytokine production (**Fig. S6e**), consistent with the known role of A20 in regulating inflammatory responses^49^.

Collectively, these data identify an NF-κB-driven regulatory circuit which constrains T-cell activation through the dynamic formation of a super-enhancer that drives expression of *TNFAIP3*, a key NF-κB inhibitor. In primary CD4 T-cells, this circuit is disrupted – and super-enhancer formation prevented – by the risk variant at rs6927172, thus revealing the biological effect of a pleiotropic disease association.

## DISCUSSION

A fundamental goal of GWAS is to better understand disease biology^8^. As such, despite widespread success in variant discovery, this goal remains largely unfulfilled – since we have not yet managed to transition from lists of associated SNPs to insights into disease mechanisms. Here, we have adapted an MPRA to simultaneously assess the functional consequences of hundreds of non-coding genetic variants in primary CD4 T cells – the cell-type whose regulatory DNA is most enriched for immune-mediated disease SNPs^2, 17^. By analysing each SNP individually, this method bypasses the limitations of LD and enables putative causal variants to be identified based on their functional consequences. Unlike fine-mapping, this approach does not attempt to refine GWAS statistics and so does not provide a specific estimate of causality for each SNP. However, by identifying SNPs that causally change biology, this method can establish the functional consequences of disease-associated genetic variation and provide a focus for studying disease mechanisms. Importantly, this approach is broadly applicable and could be used to identify putative causal variants at any disease-associated locus that overlaps with T cell regulatory elements – even those that cannot be fine-mapped – thus overcoming a major bottleneck in the transition from genetic variants to disease mechanisms.

To illustrate the value of this approach, we use the MPRA results as a basis for investigating possible disease mechanisms at a pleiotropic locus that has been linked to 6 different immune-mediated diseases, but cannot be fine-mapped. In doing so, we uncover a regulatory circuit that constrains T cell activation through the dynamic formation of *TNFAIP3* super-enhancer, and show how this can be disrupted by a common, expression-modulating variant that perturbs NF-κB binding – consistent with the known vulnerability of super-enhancers to perturbation of their components^9, 57, 58^. Altogether, this identifies a molecular and cellular mechanism that is likely to be broadly involved in human autoimmune disease. Indeed, while exuberant effector CD4 T cell responses have been implicated in the initiation and perpetuation of all of the diseases linked to rs6927172 (refs 59,60), direct evidence of how common genetic variation might contribute to this has previously been lacking.

A key strength of performing MPRA, and subsequent follow-up experiments, in primary cells is that this provides greater confidence that the results are physiologically relevant. For example, these data show that in primary CD4 T cells, the biological effect of this multi-disease-associated haplotype is solely mediated by *TNFAIP3*, and not any of the other genes at the locus. There are many lines of evidence supporting T cells as the relevant cell-type for this association, including the presence of a T cell-specific super-enhancer involving the lead SNP^9^, the enrichment of the super-enhancer components for transcription factors involved in T cell activation, and the central role that T cells play in the diseases linked to rs6927162^59, 60^. Moreover, a very recent a study of chromatin accessibility identified an allele-specific effect of rs6927172 in stimulated CD4 T cells that was not detected in other immune cell-types^36^. By resolving the downstream consequences of this effect – both on local transcription and, in turn, on T cell responses – we extend this observation and identify a mechanism consistent with a pleiotropic predisposition with to multiple diseases.

Of note, a similar mechanism was previously reported for a different locus that is located downstream of *TNFAIP3* and is associated with SLE, but not with any the other diseases linked to rs6927172 (ref 61). At this low frequency haplotype, which is not in LD with rs6927172 (r^2^ = 0.001), the putative causal variant also alters NF-κB binding and interacts with the *TNFAIP3* promoter^61^. That two distinct disease-associated loci have similar functional consequences highlights the importance of *TNFAIP3* in human autoimmunity, and may point to cell-type specific effects. For example, the SLE-only locus does not interact with *TNFAIP3* in stimulated CD4 T cells^33^ but has strong enhancer activity in transformed B cells^46^ – a cell type linked to SLE pathogenesis and enriched for SLE-associated variants^2^.

This adapted MPRA is subject to the same potential limitations of regular MPRA, in that each construct is tested in a plasmid, out of its native genomic context – reinforcing the importance of studying regions in relevant cells. Extending MPRA to other primary cell types, particularly to study disease-specific loci, will be an important next step. These data also highlight the value of considering MPRA results as hypotheses to be experimentally tested, rather than as definitive insights in isolation. Indeed, we show that using multiple biological replicates adds considerable power to identify expression-modulating effects, and so experimentally characterising the functional consequences of any result is essential.

In summary, we have developed a scalable method that is able to distil disease-associated haplotypes down to specific functional variants in relevant primary cells, thereby generating testable hypotheses into disease mechanisms – even within gene deserts – while overcoming some of the limitations of statistical fine-mapping. This can provide important insights into disease biology, and represents a generalisable framework by which the considerable potential of GWAS in immune-mediated disease could finally be realised.

## Acknowledgments

We acknowledge J.Lewandowski, M.Parkes, A.Kaser, D.Thomas and members of the Smith and Rinn labs for helpful discussion, J.Sowerby for experimental assistance, and D.Seyres for ChIP advice. pRSCgfp-hAIM2 was a gift from Emad Alnemri (Addgene #51666), CBFRE-EGFP was a gift from Nicholas Gaiano (Addgene #17705), and pOTTC407-pAAV EF1a eGFP was a gift from Brandon Harvey (Addgene #60058). We thank NIHR BioResource volunteers for their participation, and acknowledge NIHR BioResource centres, NHS Blood and Transplant, and NHS Trusts and staff for their contribution. This work was supported by the Wellcome Trust (Intermediate Clinical Fellowship to J.C.L, 105920/Z/14/Z; Senior Fellowship to C.W., WT107881), Crohn’s and Colitis UK (M2018/3), the National Institute for Health Research [Cambridge Biomedical Research Centre at the Cambridge University Hospitals NHS Foundation Trust], the Howard Hughes Medical Institute (Gilliam Fellowship to A.G.), the Engineering and Physical Sciences Research Council & GlaxoSmithKline (iCase studentship to A.H., EP/R511870/1), the Medical Research Council (MC UU 00002/4 to C.W.) and the National Institutes of Health Oxford-Cambridge Scholars Program. The views expressed are those of the authors and not necessarily those of the NIH, the NHS, the NIHR or the Department of Health and Social Care.

## Author Contributions

Conceptualization, J.L.R., and J.C.L.; Methodology, C.B., A.G., O.S.B., and J.C.L.; Software, A.G., and K.M.; Investigation, C.B., A.G., C.G., A.H., T.H., T.A., M.W.E., and J.C.L.; Writing– Original Draft, J.C.L.; Writing – Review & Editing, all authors; Funding Acquisition, J.C.L.; Supervision, C.W., K.G.C.S., J.L.R., and J.C.L.

## Competing Interests

The authors declare no competing financial interests

## METHODS

### Primers

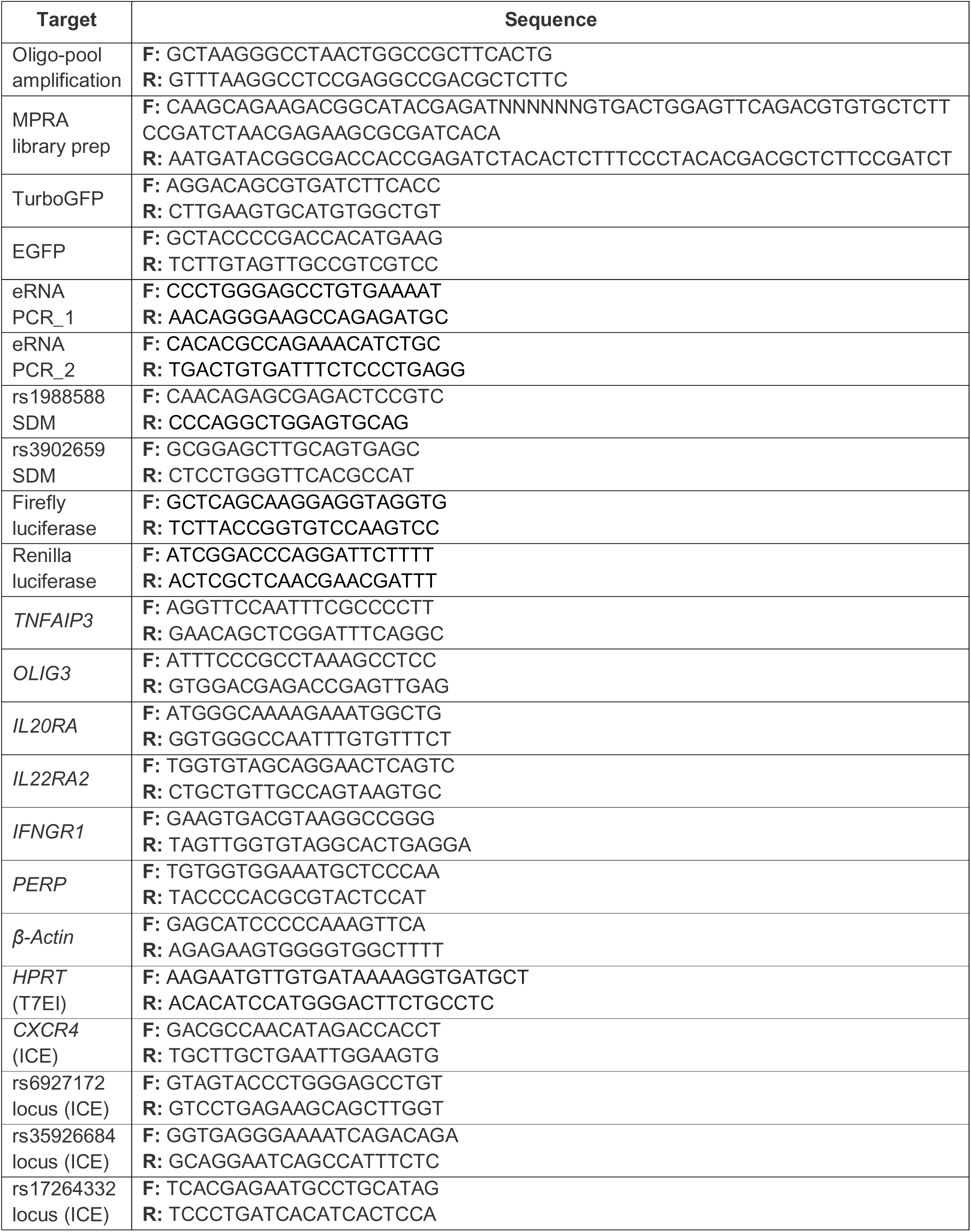
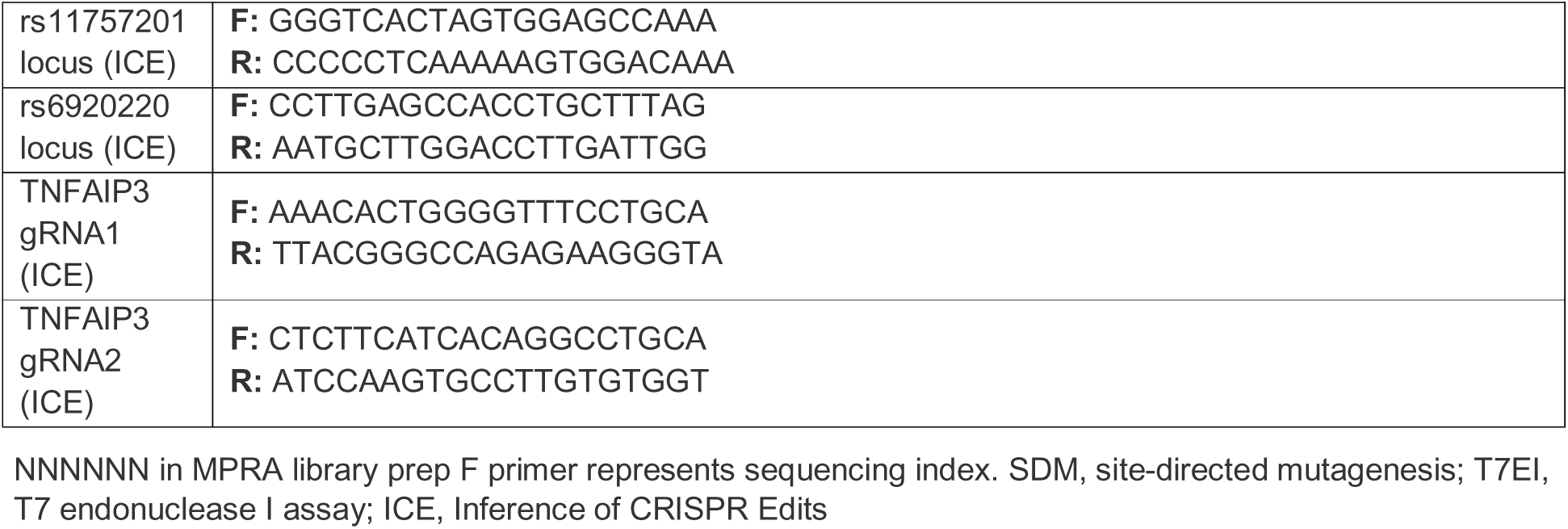

### Region selection and library design

Regions for study were identified from published immunochip studies in 10 immune-mediated diseases. Immunochip studies were used because this genotyping chip was designed to provide dense SNP coverage of established loci. Criteria for inclusion were: no coding genes or well-characterised non-coding genes within the extended haplotype tagged by all SNPs in LD with the lead variant (r^2^>0.8) and extended by 50kb at each side. 14 regions were selected (**Table 1**). For each region, oligonucleotides were designed to test the expression-modulating effect of every SNP in the associated haplotype (r^2^>0.8 with the lead SNP; total = 264 variants) and to tile the locus at 50bp intervals. Allelic constructs for each SNP were designed using 3 sliding windows around the SNP, such that 1/3, 1/2, and 2/3 of the construct were located 3’ of the variant in each construct – as has been used previously^12^. If adjacent SNPs were located within 114 bp of one another, additional oligonucleotides were synthesised to test the combination of risk alleles. Each allelic construct was tagged by 30 unique 11nt barcodes, and each tiling construct was tagged by 6 unique barcodes. Ten positive control SNPs were included: 5 were proven expression-modulating variants in lymphoblastoid cells^11^, 2 were single variant eQTLs^2^, and 3 were synthetically designed to include / disrupt a consensus binding motif for transcription factors active in CD4 T cells (GATA1/3 motif = TGATAG; RUNX1 motif = TGTGGTTT; NF-κB motif = GGGGGAATCCC). For the tiling analysis, positive and negative control regions (each 2kb) were included from T cell super-enhancers (chr21:36421330-36423329 and chr1:198626200-198628199, hg19) and gene deserts without any evidence of enhancer activity (chr4:29562525-29564524 and chr4:34780413-34782412, hg19) respectively. In total, 99,990 170bp oligonucleotides were synthesised (Twist Biosciences) to contain, in order, the 16-nt universal primer site ACTGGCCGCTTCACTG, a 114-nt variable genomic sequence, KpnI and XbaI restriction sites (TGGACCTCTAGA – for insertion of the GFP reporter cassette), an 11-nt unique barcode sequence, and the 17-nt universal primer site AGATCGGAAGAGCGTCG.

### Oligonucleotide library cloning

Oligonucleotide libraries were re-suspended in nuclease-free water and amplified by emulsion PCR (NEB Q5 polymerase, 30 cycles, Micellula DNA Emulsion & Purification Kit (Chimerx)) using primers containing SfiI restriction sites. 200ng of the purified PCR-amplified oligonucleotide library was digested with SfiI (NEB) and cloned into SfiI-digested pGL4.10M vector^10^ using One Shot MAX Efficiency DH5-T1R Competent *E.coli* (ThermoFisher). Plasmids were purified using Plasmid Plus Maxi kits (Qiagen), quantified (Nanodrop 1000 spectrophotometer, ThermoFisher) and sequenced to check library complexity. 2μg purified plasmids were digested with KpnI/XbaI (NEB) and ligated with a KpnI/XbaI-digested fragment containing a promoter and reporter gene (EGFP). Initial ligation was performed using a minimal promoter–GFP reporter cassette^66^. Subsequent ligations, to test the activity of other promoters in primary CD4 T cells, were performed using: SV40 promoter (derived from CBFRE-EGFP), RSV promoter (derived from pRSCgfp-hAIM2), and EF1α promoter (derived from pOTTC407-pAAV EF1a eGFP). In each case, ligation products were transformed into *E,coli*, purified and quantified as described above. The final pooled MPRA plasmid library was sequenced (MiSeq) to confirm sufficient oligonucleotide representation.

### CD4 T cell purification, transfection and cell culture

Source Leukocytes, freshly purified from healthy donors, were obtained from Massachusetts General Hospital (MGH) Blood Transfusion Service (BTS). Peripheral blood mononuclear cells were isolated by density centrifugation using Histopaque 1077 (Sigma), and CD4 T cells were positively selected using immunomagnetic microbeads and LS columns (Miltenyi Biotec). Purity was confirmed to be >95% by flow cytometry (data not shown). CD4 T cells were washed, counted and split 2:1 for immediate nucleofection (resting) or stimulation. Stimulation was performed for 4 days using recombinant human IL-2 (10ng/ml, Peprotech) and Anti-Biotin MACSiBead Particles loaded with CD2, CD3, and CD28 antibodies (bead-to-cell ratio 1:2, Miltenyi Biotec). Resting or stimulated CD4 T cells were nucleofected with the MPRA plasmid library using at least 6 technical replicates for each sample, which were later pooled (for each replicate: 5μg vector (in 5μl) transfected into 5 million CD4 T cells in 100μl 1M nucleofection solution^67^) using a Nucleofector 2b device (Lonza; program V024 for resting T cells and T023 for stimulated T cells). After nucleofection, 500μl pre-warmed media was immediately added to each cuvette and cells were gently transferred to a 6-well flat-bottomed plate (final volume per well = 5ml, equivalent to 1 million cells per ml) and cultured at 37°C, 5% CO_2_. Cell culture media: Iscove’s Modified Dulbecco’s Medium (ThermoFisher) containing 20% Fetal Bovine Serum (ThermoFisher), 1% non-essental amino acids (ThermoFisher), 2mM glutamine (GlutaMAX, ThermoFisher) and 1% sodium pyruvate (ThermoFisher). No antibiotics were included. 24 hours after nucleofection, cells were harvested, pooled and lysed in RLT Plus buffer (Qiagen) containing 1% 2-mercaptoethanol.

### Flow cytometry

CD4 T cell purity and composition, and transfection efficiency were assessed by flow cytometry using a BD LSR II flow cytometer (HSCRB Flow Cytometry Core Facility). Purity and composition panel: CD4 APC, CCR4 BV421, CCR6 AF700, CD3 FITC, CD62L PerCP Cy5.5, CXCR3 PE Dazzle 594, CD45RA PE/Cy7, Zombie Aqua Fixable Viability Kit (all Biolegend), Fc receptor blocking reagent (Miltenyi). Transfection efficiency panel: GFP (from MPRA plasmids), CD4 APC, Zombie Aqua Fixable Viability Kit (both Biolegend), Fc receptor blocking reagent (Miltenyi). Data were gated using FlowJo v10 (BD).

### Library preparation

Lysates were DNA depleted using a gDNA eliminator column (Qiagen) and RNA was extracted using a RNeasy Plus micro kit according to the manufacturer’s instructions (Qiagen). For library preparation, 1μg RNA was treated with TURBO DNAse (ThermoFisher) and reverse transcribed (SuperScript IV VILO, ThermoFisher) according to the manufacturer’s instructions. DNA removal was confirmed by qPCR for EGFP and compared to a no-RT control (all performed in triplicate). Sequencing libraries were prepared by PCR amplification (30 cycles, annealing temperature 55C) using PfuUltra II Fusion DNA polymerase (Agilent) and custom primers that were designed to anneal to a 3’ site within the EGFP gene (F) and the 3’ universal primer site within the oligonucleotide sequences (R). These primers contained sequencing indices to enable multiplexing. Amplified libraries were cleaned using sequential SPRI bead clean-up (0.6X, 1.6X, 1.0X; Agencourt AMPure XP, Beckman Coulter). Four sequencing libraries were made from the input MPRA plasmid library using 50ng vector and 18 PCR cycles (other conditions were the same as for the RNA libraries). The quality and molarity of all libraries was assessed using a BioAnalyzer 2100 (Agilent) and the libraries were sequenced in pools of 6 (Illumina HiSeq2500 high output flow-cell, 50bp, single-end reads) – median 39.7 million reads per sample.

### MPRA analysis

#### Pre-processing

Barcode counts were obtained from the FASTQ files for each sample after quality control (FastQC). To be counted, a sequenced barcode had to be a perfect match for an oligonucleotide library barcode and be followed by at least 10 bases of the expected constant sequence (XbaI restriction site and GFP). To be deemed a successful transfection, at least 70% of the oligonucleotide library had to be represented in the resulting count file (i.e. barcode count ≥1). Raw count data were then normalised to correct for sequencing depth (counts per million mapped reads, cpm) and then filtered to remove barcodes with a median cpm < 0.5 in either the RNA or DNA samples (equivalent to a raw barcode count of ∼20-25 reads). Barcode counts for identical constructs were then collapsed (summed) and quantile normalised.

#### Principal component analysis

Principal component analysis (PCA) was performed on pre-processed construct-level barcode counts from RNA samples using the *prcomp* function in the *stats* package in R (version 3.5.1). In brief, the pre-processed data were centered and scaled to have unit variance and then singular value decomposition was performed on the resulting data matrix. No components were omitted. The first two components in the resulting data object are plotted in Figure 1C. Eigenvalues representing the total amount of variance in the data explained by each component are shown.

#### Pairwise correlation analysis

For every sample, the transcriptional activity of each construct was calculated by dividing the normalised construct-level barcode count (mRNA) by the median normalised count for the same construct from four sequencing replicates of the input plasmid library (DNA). Correlation matrices were separately created for resting and stimulated CD4 T cell samples using the *cor* function (Pearson correlation) in the *stats* package in R (version 3.5.1). The *reshape* package was then used to melt each correlation matrix, and these were plotted using the *ggplot2* package in R.

#### Tiling analysis

To assess for enhancer activity within each disease-associated locus, we used the *sharpr2* package^68^ in R (version 3.5.1). This was used because: (1) the tiled regions were of different sizes, (2) the offset between constructs (50bp) was not a factor of the length of genomic sequence (114bp), (3) this method facilitated inclusion of the reference allele constructs from SNPs to improve coverage within the locus (since these constructs also contained the reference genomic sequence at the sites of SNPs), and (4) none of the co-ordinates of the regions on their respective chromosomes overlapped. After subsetting the pre-processed construct-level barcode counts to remove alternate allele (SNP) constructs, the median counts for the remaining constructs in RNA and DNA samples and their genomic co-ordinates were used for analysis. The *sharpr2* function was used with default settings and without filtering on size or fragment count since the sizes were identical and the data were already filtered. The regulatory scores for each region were based on standardized log(RNA/PLASMID) and a regional FWER cutoff (0.05) was used to call to call high resolution driver elements indicative of enhancer activity.

#### SNP analysis

For the SNP analysis, we used QuASAR-MPRA^32^, implemented in the *QuASAR* package in R, since this accounts for potential uncertainty in the original plasmid proportions, over-dispersion, and sequencing errors. After pre-processing to normalised construct-level barcode counts, and removing enhancer constructs, 964/970 SNP constructs were available for analysis. For every SNP construct, the numbers of reference allele reads and alternate allele reads in each RNA sample, and the proportion of reference allele reads in the DNA vector were used as input. This was implemented using the *fitQuasarMpra* function, and uses a beta-binomial distribution to model the imbalance in the allelic constructs, since this better calibrates the P values under the null hypothesis than other methods. Since the *fitQuasarMpra* function can only analyse one sample at a time, a standard fixed-effect meta-analysis was used to combine the results for each SNP construct for the biological replicates – as recommended by the authors of this method. The *fitQuasarMpra* function provides the logit transformation of the proportion of reference reads in RNA (*β*_l_) and the standard error of this estimate 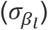. We then calculated logit transformation of the proportion of reference reads in DNA (*β*_0_) in order to perform the meta-analysis for *k* samples as follows:

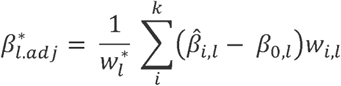

where 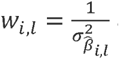 and 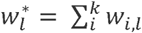. We then calculated a meta-analysis Z score and P value:

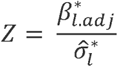

where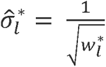.

Correction for multiple testing was performed by controlling the false discovery rate^69^.

### Luciferase-based validation

Geneblocks corresponding to the genomic sequences of the reference and alternate alleles of the lead expression-modulating SNPs at each haplotype were synthesised with flanking restriction sites (5’ KpnI, 3’ BamHI; IDT). For one haplotype (14q32 – associated with T1D) the sequence of the lead SNP construct was too GC rich to be synthesised and so the corresponding region was PCR amplified from a major allele homozygote and site-directed mutagenesis was used to create the alternate allele constructs (Q5 Site-Directed Mutagenesis Kit, NEB – used according to manufacturer’s instructions). Additional geneblocks for 2 positive control SNPs and 5 SNPs with no expression-modulating activity were also synthesised (IDT). Geneblocks were then KpnI/BamHI-digested and ligated into a similarly digested custom Firefly luciferase vector (synthesised by VectorBuilder) such that they were inserted immediately proximal to the Firefly luciferase promoter (EF1α). The ligation product was transformed into *E.coli*, sequenced to confirm successful insertion (Genewiz) and purified and quantified as described above. For each geneblock, equimolar amounts of the Firefly vector and a custom-designed Renilla luciferase vector (total 5μg vector mix in 5μl water) were nucleofected into resting CD4 T cells (program V024, Nucleofector 2b). After 24 hours, cells were harvested, lysed and DNA and RNA were extracted from the lysate using the AllPrep DNA/RNA Micro kit (Qiagen) and quantified (Nanodrop 1000 spectrophotometer, ThermoFisher). 200ng RNA was DNAse treated (TURBO DNase, ThermoFisher) and reverse transcribed (SuperScript IV VILO, ThermoFisher). Quantification of Firefly and Luciferase genes in extracted DNA and mRNA (cDNA) samples was performed in triplicate using qPCR. The results were normalised using an adaptation of the Delta-Delta-Ct method in which the Cts for Firefly and Renilla (mRNA) were first normalised to their respective DNA Cts (to control for any imbalance in the transfected vector mix) and then each Firefly delta-Ct was normalised to the Renilla Delta-Ct (to control for transfection efficiency). This produced a measure of the activity of each allelic construct, which was compared between the reference and alternate alleles at each SNP to provide an estimate of the expression-modulating effect. Four biological replicates were performed. For each SNP, the expression-modulating effects observed in the MPRA and validation experiments were plotted, and linear regression was performed using the *lm* function in the *stats* package in R (version 3.5.1).

### Fine-mapping ankylosing spondylitis association on 2p15

The ankylosing spondylitis summary statistics^21^ were downloaded from the GWAS catalog and SNPs in the region chr2:62518445..62618445 (build hg19) were extracted. Using an established approach^70^, approximate Bayes factors summarising the association at each SNP, and thus the posterior probabilities for each SNP to be causal, were calculated^71^ – assuming a single causal variant in the region. These posterior probabilities were used to construct a 99% credible set, which contained 4 SNPs and was expected to contain the true causal variant with 99% probability. Recent work has shown that this conventional procedure can be biased, but that any such bias can be corrected^72^. We therefore used the corrcoverage R package to correct any bias^72^ and identified a 99% credible set containing three SNPs: rs6759298, rs4672505 and rs13001372, which has a corrected coverage estimate of containing the true causal variant of 99.2%.

### NF-κB binding site analysis

A common NF-κB binding motif that can bind to all NF-κB dimers was identified from publicly available protein-binding microarray data, using a complementary approach to that described by the authors of the paper (Additional File 2 from ref 39). In brief, the reported z-scores for the affinity of 9 NF-κB dimers for each 11-mer sequence on the protein-binding microarray were combined using Stouffer’s method and the combined z-score was used to calculate the statistical significance of the overall binding. After correcting for multiple-testing using the Bonferroni method, 100 statistically significant 11-mer sequences were identified (P_adjust_ < 0.05) which had positive z-scores for every dimer. These sequences were used to generate a common NF-κB binding motif logo using Weblogo^73^.

### NF-κB immunoprecipitation following MPRA library nucleofection

CD4 T cells were purified and immediately nucleofected with the MPRA plasmid library as described earlier (n = 4). After 24 hours, cells were harvested, counted and resuspended in fresh cell culture media (10^6^ cells/ml). For cross-linking, 37% Formaldehyde was added to a final concentration of 1%, and cells were placed on a rocker for 10 minutes (room temperature). Cross-linking was quenched by adding Glycine (final concentration 0.125M) and shaking for 5 min (room temperature). Cells were washed twice in ice cold PBS and cell pellets were lysed for 10 min at a density of 10^7^ cells/ml in lysis buffer supplemented with Protease inhibitor (Complete Mini EDTA-free Protease Inhibitor cocktail tablets; Roche). Lysis buffer: 50mM HEPES pH 7.9, 140mM NaCl, 1mM EDTA pH 8.0, 10% v/v Glycerol, 0.5% v/v IGEPAL CA-630, 0.25% v/v Triton X-100. 2 cycles of sonication (30s ON/30s OFF) were performed using a Bioruptor Pico (Diagenode) to remove contaminants while minimising chromatin shearing^74^. Triton X-100 and NaCl were added to a final concentration of 1% and 100mM, respectively, and the samples were frozen at −80°C until further use. 10µg of sheared chromatin were cleared by centrifugation (21,000G, 10min, 4°C) and immunoprecipitation was performed overnight at 4°C using an anti-NFkBp65 antibody (clone D14E12; Cell Signaling Technology 8242) or an isotype control (rabbit IgG monoclonal antibody, abcam; ab172730) with the SimpleChIP Plus Sonication ChIP kit (Cell Signaling Technology) – according to the manufacturer’s instructions. Sequencing libraries were prepared from isolated plasmids as described above (26 PCR cycles) and sequencing was performed as for MPRA libraries.

### Allele-specific NF-κB ChIP

A fresh 100ml blood sample was obtained from 8 healthy individuals who were heterozygous at rs6927172 – identified through the NIHR BioResource, a genotype-recallable panel of over 20,000 individuals. All participants provided written informed consent and ethical approval was provided through the Cambridgeshire Regional Ethics Committee (REC:08/H0308/176). CD4 T cells were purified as described earlier and left resting or stimulated for 4 days in complete RPMI supplemented with 10ng/ml recombinant IL-2 (complete RPMI: RPMI-1640 containing 10% Fetal Bovine Serum, 2mM glutamine, 10mM HEPES, 1% sodium pyruvate, 50μM 2-Mercaptoethanol, and Penicillin-Streptomycin 100u/ml (ThermoFisher)). Stimulation was performed using Anti-Biotin MACSiBead Particles loaded with CD2, CD3, and CD28 antibodies (Miltenyi Biotec) as described earlier. After 4 days, cells were harvested, cross-linked, quenched and lysed as described earlier. Lysates were washed twice with wash buffer (10mM Tris-HCl pH 8.0, 200mM NaCl, 1mM EDTA pH 8.0, 0.5mM EGTA pH 8.0) and nuclei were prepared by washing twice with shearing buffer (0.1% w/v SDS, 1mM EDTA, 10mM Tris-HCl pH 8.0). Nuclei were resuspended in 200µl shearing buffer per 10^7^ cells and sonicated for 9 cycles (30s ON/30s OFF) using a Bioruptor Pico. Triton X-100 and NaCl were added to the sheared chromatin to a final concentration of 1% and 100mM, respectively. The sheared chromatin was frozen at −80°C until further use. NF-κB ChIP was performed as described above. To assess for allele-specific binding, the NF-κB-bound DNA was genotyped in triplicate (TaqMan genotyping assay C___1575580_10) alongside pre-mixed DNA from a minor and a major allele homozygote at rs6927172. A series of different ratios of minor to major allele homozygote DNA were used (from 4:1 to 1:4) to create a standard curve, against which the ratio of FAM:VIC intensities in the NF-κB ChIP samples were compared.

### Allele-specific eRNA analysis

DNA and RNA were extracted (AllPrep DNA/RNA kit, Qiagen) from stimulated CD4 T cell lysates (5 x 10^6^ cells, un-nucleofected) that were stored as part of the MPRA experiment. Genotyping was performed to identify 6 heterozygotes at rs6927172 (TaqMan genotyping assay C___1575580_10). RNA was TURBO DNase-treated and reverse-transcribed as described earlier. Nested PCR was performed to amplify the region surrounding rs6927172 from genomic DNA and cDNA. PCR amplicons were gel-purified (Zymoclean Gel DNA recovery kit, Zymo), quantified (Nanodrop 1000 spectrophotometer) and diluted to 8ng/μl. 1μl (equivalent to a 5:1 insert:vector ratio) was ligated into a blunt-ended TOPO vector (Zero Blunt TOPO PCR Cloning Kit, ThermoFisher) and transformed into *E.coli*, according to the manufacturer’s instructions. For each sample, 96 colonies were picked and genotyped to measure allelic ratios (TaqMan genotyping assay C___1575580_10).

### H3K27ac ChIP-seq and analysis

A fresh 100ml blood sample was obtained from 3 major allele homozygotes and 3 minor allele homozygotes at rs6927172. All were identified via the NIHR BioResource. CD4 T cells were purified and stimulated for 4 days using anti-Biotin MACSiBead Particles loaded with CD2, CD3, and CD28 antibodies (Miltenyi Biotec) and 10ng/ml recombinant IL-2 in complete RPMI, as described earlier. After 4 days, cells were harvested, cross-linked, quenched, lysed, washed, and nuclei prepared and sheared as described earlier. 2% input samples were stored prior to immunoprecipitation. Immunoprecipitation was performed overnight at 4°C with rotation using an anti-H3K27ac antibody (abcam; ab4729) or an isotype control (rabbit IgG monoclonal antibody, abcam; ab172730) with the SimpleChIP Plus Sonication ChIP kit (Cell Signaling Technology) – according to the manufacturer’s instructions. 50ng of immunoprecipitated DNA or input sample were used to prepare sequencing libraries using the iDeal Library Preparation kit (Diagenode), according to manufacturer instructions. 10 PCR cycles were used for amplification. The quality and molarity of all libraries was assessed using a BioAnalyzer 2100 (Agilent) and the libraries were sequenced in pools of 8, with each pool being sequenced in 2 lanes of an Illumina HiSeq2500 high output flow-cell (50bp, single-end reads) – median 50.4 million total reads per H3K27ac sample and 79.6 million total reads per input sample. Sequencing reads were trimmed to remove low quality base calls and residual adaptors at the 3’ end using TrimGalore! (Phred score 24) and filtered to remove reads shorter than 36bp https://w.bioinforcs.babraha.ac.uk/projects/trim_galore/. Trimmed reads were then aligned to the reference human genome (hg19) using Burrows-Wheeler Aligner (BWA) with default parameters^75^. Aligned reads were converted to BAM files, sorted, and technical duplicates merged before indexing – all using SAMtools^76^. PCR duplicates were identified using Picard tools (http://broadinstitute.github.io/picard/) and removed together with unmapped reads using SAMtools. The resulting BAM files were re-sorted and indexed after filtering. For visualisation in IGV ^77^, bigwig files were generated using bamCoverage (deepTools2, ref 78, with the following parameters: --binSize 5 --normalizeUsing CPM --effectiveGenomeSize 2685511504 --extendReads 200. Biological replicates and inputs for each genotype were then merged using SAMtools. Peaks were identified in H3K27ac and input libraries using MACS2^79^ after downsampling the input files to have the same number of reads as the H3K27ac samples. The following parameters were used: -g hs -f AUTO --qvalue 0.01 -B --nomodel --extsize=200. MACS peaks of H3K27ac were used as constituent enhancers for super-enhancer identification using Rank Ordering of Super-Enhancers (ROSE; https://bitbucket.org/young_computation/rose)^46^. A stitching distance of 12,500bp and a promoter exclusion zone ± 2,000bp were used. For the minor allele homozygote samples, the activity at the super-enhancer locus was calculated (for comparison with the major allele homozygotes) by summing the activity of all detected enhancers within the region and normalising for region size.

### *In silico* transcription factor binding analysis

Transcription factor binding motifs that were enriched at constituent elements within the 6q23 super-enhancer were identified using TRAP (multiple sequences)^80^, with all human promoters as the reference dataset and a Benjamini-Hochberg FDR correction for multiple testing^69^. Motifs were obtained from the Jaspar CORE vertebrate database. Pathway analysis of enriched transcription factors within annotated KEGG pathways was performed using g:Profiler (https://biit.cs.ut.ee/gprofiler/gost) with a Benjamini-Hochberg FDR correction for multiple testing^69^.

### Promoter-capture Hi-C analysis

Interactions of the 6q23 super-enhancer in stimulated CD4 T cells were identified from an existing promoter-capture Hi-C dataset^33^ using the capture Hi-C plotter (https://www.chicp.org). Genetic association data for the 6q23 locus was based on IBD summary statistics^62^.

### qPCR in CD4 T cells from IBD patients

131 patients with active ulcerative colitis or Crohn’s disease were recruited before commencing treatment as part of in a separate study^81, 82^. All patients provided written informed consent and ethical approval was provided by the Cambridgeshire Regional Ethics committee (REC:08/H0306/21). CD4 T cells were positively selected from a fresh 100ml blood sample using immunomagnetic microbeads, as described earlier. Cells were immediately lysed and RNA and DNA were subsequently extracted using an AllPrep DNA/RNA Mini kit (Qiagen). Genotyping was performed using the Illumina Human OmniExpress12v1.0 BeadChip, according to the manufacturer’s instructions, and data were processed as previously described^83^. qPCR for genes at the 6q23 locus was performed in triplicate, using custom exon-spanning primers, with beta-actin as a reference gene (QuantiFast SYBR Green PCR Kit; Qiagen) on a Roche LightCycler 480. All primers were first validated by amplicon Sanger sequencing.

### Guide RNAs

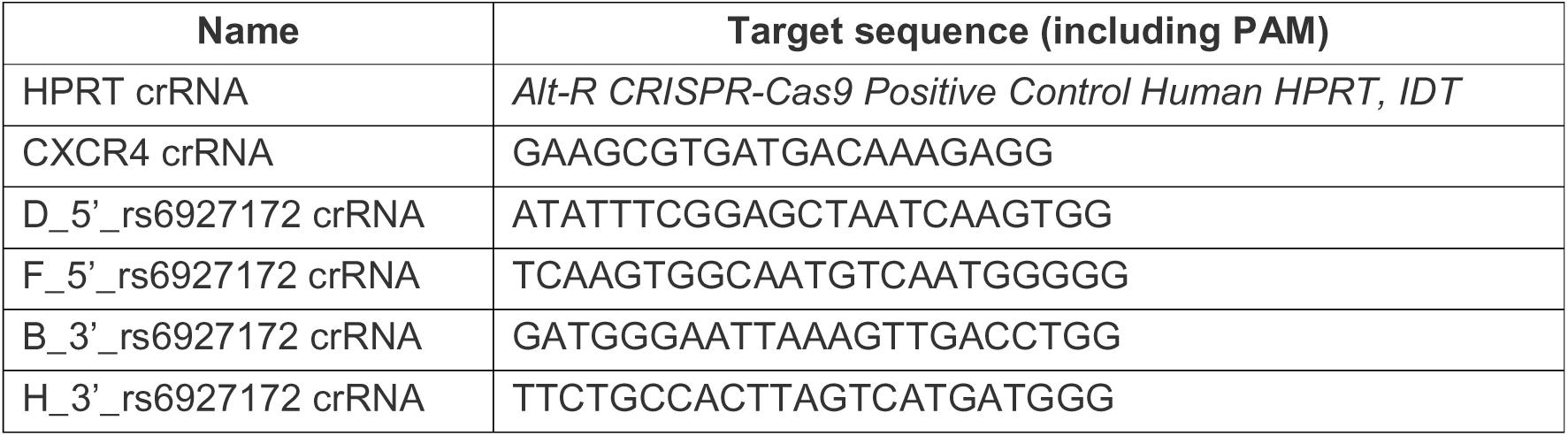
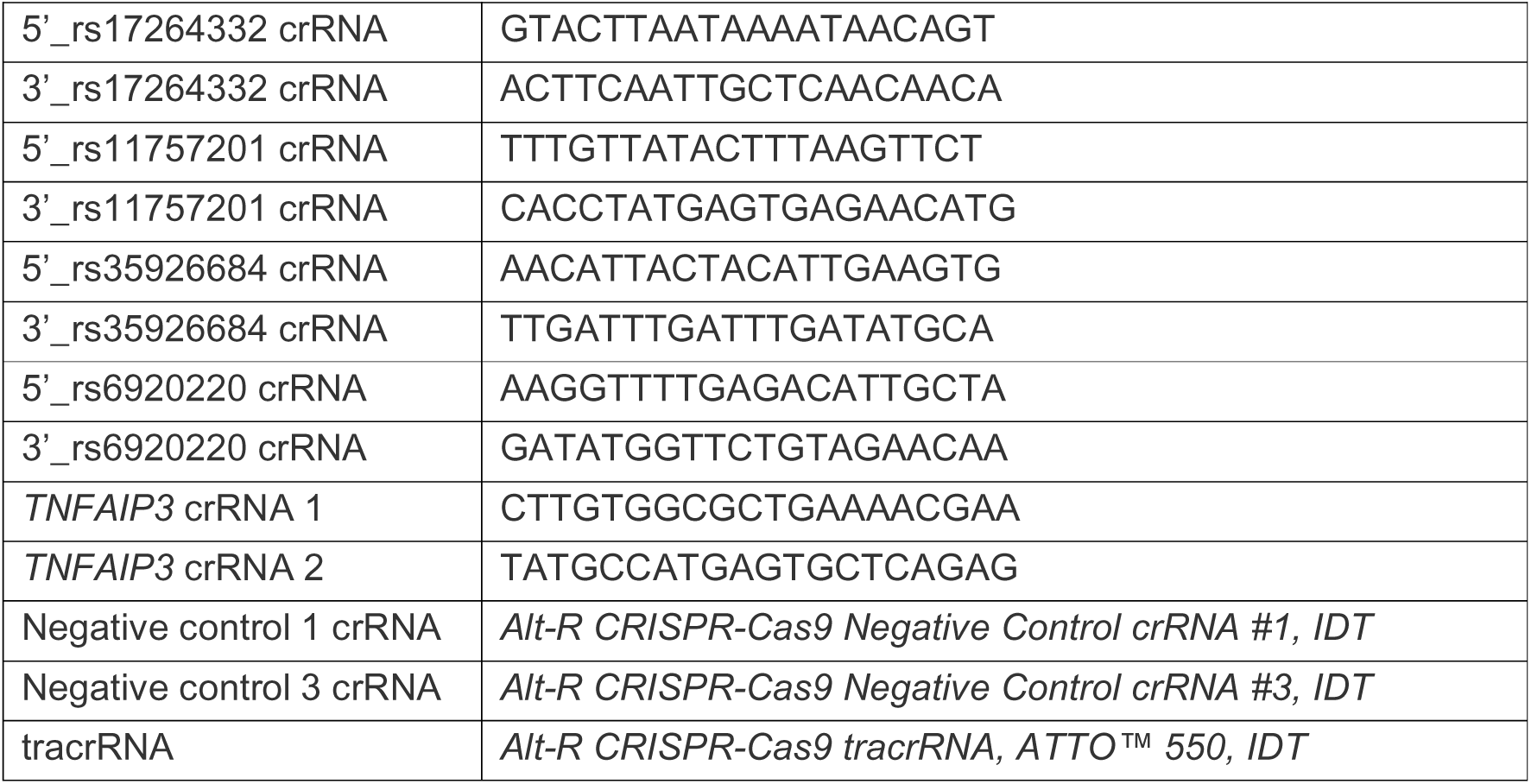

### CRISPR-Cas9 editing in resting CD4 T cells

#### Optimisation

A series of conditions were tested to find a suitable method for ribonucleoprotein (RNP)-based CRISPR editing in resting CD4 T cells, including different nucleofection buffers (Human T cell Nucleofector kit, Lonza; 1M nucleofection solution^67^), nucleofection programs (U014; V024), and use or not of an electroporation enhancer (Alt-R Cas9 Electroporation Enhancer, IDT). All nucleofections were performed using 10^6^ freshly-purified CD4 T cells in 100 μl nucleofection buffer using a Nucleofector 2b device (Lonza), based on previous optimisation studies (data not shown). CD4 T cells were positively selected from peripheral blood mononuclear cells isolated from fresh single leukocyte cones (National Blood Service, Cambridge, UK) as described earlier. A series of control gRNAs were used – each of which was synthesised as a crRNA and combined with a tracRNA to form a functional gRNA duplex. Positive control gRNA targets: *HPRT* (Alt-R CRISPR-Cas9 Positive Control crRNA, Human HPRT; IDT), *CXCR4* (ref 53). Negative (non-targeting) controls (NTC): Alt-R CRISPR-Cas9 Negative Control crRNA #1 and #3 (IDT). crRNAs and tracrRNA (Alt-R CRISPR-Cas9 tracrRNA, ATTO™ 550) were synthesised by IDT, and reconstituted in duplex buffer at 200μM. gRNA duplexes were generated by mixing 200μM tracrRNA with 200μM crRNA in a 1:1 ratio, and heating the mix to 95°C for 10 minutes before slowly cooling to room temperature. Any unused gRNA duplex was stored at −80°C. Cas9 RNPs were generated immediately before use by adding high fidelity Cas9 (Alt-R S.p. HiFi Cas9 Nuclease V3, 61μM; IDT) to the gRNA duplex in a 1:3 ratio, and incubating the mix at 37°C for 20 min – producing 15μM Cas9 RNP. 5μl Cas9 RNP (containing ∼18μg Cas9) was then nucleofected into CD4 T cells. The electroporation enhancer was reconstituted in nuclease-free water to a concentration of 400μM, and 1μl was added to the Cas9 RNP where indicated (equivalent to a final concentration of ∼4μM in the nucleofection reaction). After nucleofection, 500μl pre-warmed media was immediately added to the cuvette and cells were gently transferred to a 24-well flat-bottomed plate (final volume per well = 1ml) and cultured at 37°C, 5% CO_2_. Cell culture media: X-VIVO15 (STEMCELL) supplemented with 5% FBS, 50μM 2-mercaptoethanol, and 10μM N-acetyl l-cystine.^84^ After 6 hours the media was changed to optimise viability, and fresh pre-warmed media containing low dose recombinant human IL-7 (1ng/ml; Peprotech) was added to promote T cell survival without stimulation^85^. 48 hours after nucleofection, the media was changed for pre-warmed media containing anti-Biotin MACSiBead Particles loaded with CD2, CD3, and CD28 antibodies (bead-to-cell ratio 1:2, Miltenyi Biotec) and IL-2 (10ng/ml) in order to stimulate T cells. Cells were harvested 24 hours after stimulation and either used for flow cytometry or lysed in RLT Plus buffer containing 1% 2-mercaptoethanol. Surface expression of CXCR4 was assessed in edited and non-targeting control cells by flow cytometry: CXCR4 APC, Zombie Aqua Fixable Viability Kit (all Biolegend), Fc receptor blocking reagent (Miltenyi). Viability was ∼80% (of total cells) at the end of the experiment.

#### Editing efficiency assessment

DNA/RNA extraction was performed from cell lysates using the AllPrep DNA/RNA Micro Kit (Qiagen). For initial optimisation experiments using an *HPRT* gRNA-containing RNP (or non-targeting control), editing efficiency was estimated using a T7 Endonuclease assay (Alt-R Genome Editing Detection Kit, IDT). In brief, 50ng DNA from the positive or negative control samples were PCR amplified (30 cycles) using primers that eccentrically flanked the predicted cut-site (Phusion High Fidelity Polymerase, ThermoFisher). T7 endonuclease I digestion was then performed according to the manufacturer’s instructions. 10μg Proteinase K was added (1μl of 10mg/ml stock) to inactivate the T7 endonuclease I before fragment analysis. Digested heteroduplexes were quantified using a high-sensitivity DNA chip on a Bioanalyser 2100 (Agilent) – undigested band: 1050-1600bp; digested bands: 250-300bp and 700-1000bp. The optimal conditions for RNP nucleofection in resting CD4 T cells were identified as: 1M nucleofection buffer, V024 program, with electroporation enhancer. These conditions were used for all subsequent experiments. Editing efficiency for subsequent CRISPR experiments was estimated using ICE (Inference of CRISPR Edits, Synthego)^63^ after observing that the results correlated well with colony-based amplicon sequencing methods (data not shown). In brief, the target sequence in edited cells and non-targeting control cells was PCR-amplified, gel purified and Sanger sequenced. Sequencing traces (ab1 files) were uploaded to the ICE website (https://ice.synthego.com/<u>#/</u>) and non-negative least squares regression was used to infer the composition of indels based on the traces and the gRNA sequences, using the non-targeting control as a reference.

#### Deletion of NF-κB binding site at rs6927172 locus

gRNAs flanking the NF-κB binding site, which is disrupted by rs6927172, were designed using an online gRNA design tool (http://crispr.mit.edu) with 250bp of genomic sequence centred on rs6927172 as the target (chr6:138002050-138002300, hg19). 2 gRNAs proximal (5’) to the NF-κB motif (termed D and F) and 2 distal (3’) gRNAs (termed B and H) were selected and synthesised (IDT). These gRNAs were additionally checked for suitable on- and off-target activity with the GPP sgRNA design tool (https://portals.broadinstitute.org/gpp/public/analysis-tools/sgrna-design) which uses the “Rule Set 2” method for assessing on-target activity^86^ and the Cutting Frequency Determination to assess off-target activity. To further reduce the possibility that any observed phenotype might be due to off-target activity, and to maximise the disruption of the NF-κB binding motif, RNPs were used in combination (one 5’ gRNA-containing RNP with one 3’ gRNA-containing RNP). Predicted indels were as follows: DB, 33bp indel; DH, 50bp indel; FB, 18bp indel; FH, 35bp indel. Cas9 RNPs for each gRNA were generated as described earlier. For nucleofections, 2.5μl 5’ gRNA-containing RNP and 2.5μl 3’ gRNA-containing RNP were mixed and 1μl electroporation enhancer was added, as described earlier. For non-targeting control RNPs, 5μl Cas9 RNP was mixed with 1μl electroporation enhancer. Nucleofection of RNPs into resting CD4 T cells (positively selected from fresh single leukocyte cones as described earlier) were performed using optimised conditions and fresh media containing low dose recombinant human IL-7 was added after 6 hours (described earlier). Cells were then rested for 48 hours after nucleofection. For nascent RNA capture experiments, 5-ethynyl uridine (EU, Click-iT™ Nascent RNA Capture Kit, ThermoFisher Scientific) was added to the cell culture media at the time of the stimulation (final concentration 0.4mM). 24 hours after stimulation, the supernatant was removed and frozen for cytokine analysis, and the cells were harvested and either used for flow cytometry or lysed in RLT Plus buffer containing 1% 2-mercaptoethanol. 6 biological replicates were performed, although the RNP combination (FB) that resulted in poor editing – due to presumed steric hindrance between Cas9 molecules – was not repeated after the first two replicates due to poor editing efficiency.

#### Deletion of other candidate SNPs in the 6q23 super-enhancer

gRNAs flanking other candidate SNPs that were located within the co-ordinates of the 6q23 super-enhancer were designed using the same method as for the rs6927172 locus. Due to nucleosome positioning, a larger genomic sequence (500bp) from which to select guides was required for rs11757201. crRNAs were synthesised (IDT) and duplexed with tracrRNAs, before incorporation into gRNA-Cas9 RNPs (as described earlier). An equimolar mix of 5’ and 3’ RNPs were nucleofected into resting CD4 T cells, which were left unstimulated for 48 hours and then stimulated with anti-CD2/ CD3/CD28 microbeads and IL-2 (as described earlier). EU was added at the time of stimulation. After 24 hours, cells were harvested and DNA and RNA were extracted.

#### Editing of TNFAIP3

Two gRNA sequences targeting *TNFAIP3* were obtained from a genome-wide CRISPR screen gRNA library (Brunello^86^) and synthesised as crRNAs (IDT). These were duplexed with tracrRNAs and incorporated into gRNA-Cas9 RNPs immediately before use, as described earlier. Nucleofection of each RNP into resting CD4 T cells (positively selected from fresh single leukocyte cones as described earlier) was performed using optimised conditions and fresh media containing low dose recombinant human IL-7 was added after 6 hours (described earlier). After 48 hours, cells were stimulated with anti-CD2/ CD3/CD28 microbeads and IL-2 (as described earlier). 24 hours after stimulation, the supernatant was removed and frozen for cytokine analysis, and the cells were harvested for flow cytometry.

#### Flow cytometry

Cell surface staining was performed using: CD69 BV421 antibody, Zombie Aqua Fixable Viability Kit (both Biolegend), Fc receptor blocking reagent (Miltenyi). ATTO-550 staining (from the ATTO-550-conjugated tracrRNA) was used to distinguish cells containing the RNP from those that did not. Intracellular staining (for experiments in which the NF-κB binding site containing rs6927172 was deleted) was performed using the eBioscience Foxp3 / Transcription Factor Staining kit (ThermoFisher) and a Phospho-IkB alpha (Ser32, Ser36) eFluor 660 antibody (ThermoFisher) with an FMO (fluorescence-minus-one) control.

#### Nascent RNA capture

Following RNA extraction, EU-labelled RNA was biotinylated, precipitated overnight and purified using Dynabeads Streptavidin T1 magnetic beads, according to the Click-iT Nascent RNA Capture Kit protocol (Life Technologies). Reverse transcription was performed using bead-bound RNA (SuperScript VILO cDNA synthesis kit) and qPCR was performed in triplicate (QuantiFast SYBR Green PCR Kit; Qiagen) on a Roche LightCycler 480 using beta-actin as reference gene. Expression of target genes was then normalised to the expression level detected in a non-targeting control (NTC).

#### Cytokine quantification

For experiments in which the NF-κB binding site containing rs6927172 was deleted, T cell-derived cytokines in the cell culture supernatants were quantified in duplicate using electrochemiluminescence (according to the manufacturer’s instructions; MesoScale Discovery Immunoassay). For experiments in which TNFAIP3 was directly edited, T cell-derived cytokines were quantified in triplicate using Quantikine ELISAs (according to the manufacturer’s instructions; R&D).

## Statistical methods

Statistical methods used in MPRA analysis are described in the relevant section. For other analyses, comparison of continuous variables between two groups was performed using a paired *t*-test or one sample *t*-test when comparing against a hypothetical value. Two-tailed tests were used as standard unless a specific hypothesis was being tested. The alpha value was 0.05, and corrected for multiple-testing where indicated.

## Data availability

Raw and processed sequencing data have been deposited in GEO and are available under the following accession numbers: GSE135925 (MPRA data), and GSE136092 (ChIP data).

