## Supplementary Table 1 and Supplementary Figures for "Resolving mechanisms of immune-mediated disease in primary CD4 T cells"

| **Chr.** | **Haplotype  co-ordinates (hg19)** | **Associated disease(s)** | **Driver elements in resting CD4  T cells** | **Driver elements in stimulated CD4 T cells** | **Driver elements (bp)** |
| --- | --- | --- | --- | --- | --- |
| 1p34 | 1:38614767-38644961 | RA | 1 | 1 | 294 |
| 2p15 | 2:62551372-62585241 | AS, Ps | 1 | 1 | 557 |
| 2p24 | 2:12632889-12648793 | ATD | 1 | 1 | 310 |
| 3p24 | 3:28068294-28079185 | MS | 1 | 1 | 121 |
| 4q32 | 4:166558881-166575539 | T1D | 1 | 1 | 336 |
| 5p13 | 5:40522112-40619865 | AS | 1 | 1 | 766 |
| 6p23 | 6:14711861-14734441 | UC, CD, MS | 1 | 1 | 559 |
| 6q23 | 6:137959135-138006604 | RA, CeD, UC, CD, SLE, T1D | 1 | 1 | 479 |
| 8q24 | 8:130602181-130624205 | UC, CD | 1 | 1 | 293 |
| 8q24 | 8:128187774-128207238 | MS | 1 | 0 | 32 |
| 11q21 | 11:95311160-95320908 | ATD, vitiligo | 1 | 1 | 49 |
| 14q32 | 14:98485011-98499051 | T1D | 1 | 1 | 261 |
| 21q21 | 21:16804230-16828335 | UC, CD | 1 | 1 | 212 |
| 21q22 | 21:40463183-40468938 | AS, PSC, UC, CD | 1 | 1 | 82 |
| 21q22 | 21:36421330-36423329 | Positive control 1 | 0 | 1 | 49 |
| 1q31 | 1:198626200-198628199 | Positive control 2 | 1 | 1 | 81 |
| 4p15 | 4:29562525-29564524 | Negative control 1 | 0 | 0 | 0 |
| 4p15 | 4:34780413-34782412 | Negative control 2 | 0 | 0 | 0 |

**Supplementary table 1. Summary of tiling analysis.**

Summary results from tiling analysis in resting and stimulated CD4 T cells using the *sharpr2* package

Filled green boxes indicate the presence of high-resolution driver elements with significant regulatory activity (FWER P < 0.05) within the genomic sequence of the disease-associated haplotype. Filled grey boxes indicate that no high-resolution driver elements were identified in the region.

Driver elements (bp) indicates the total number of bases within the disease-associated region that were identified as high-resolution driver elements in either resting or stimulated T cells.

Chr., chromosome; AS, Ankylosing Spondylitis; Ps, Psoriasis; PSC, Primary Sclerosing Cholangitis; UC, ulcerative colitis; CD, Crohn’s disease; RA, rheumatoid arthritis; CeD, coeliac disease; SLE, Systemic Lupus Erythematosus; T1D, Type 1 Diabetes; ATD, autoimmune thyroid disease; MS, multiple sclerosis.

**Supplementary Table 2. Summary of MPRA analysis for putative causal SNPs from 14 autoimmune disease-associated loci in resting CD4 T cells.**

*Provided separately as .xlsx file*

**Supplementary Table 3. Summary of MPRA analysis for putative causal SNPs from 14 autoimmune disease-associated loci in stimulated CD4 T cells.**

*Provided separately as .xlsx file*

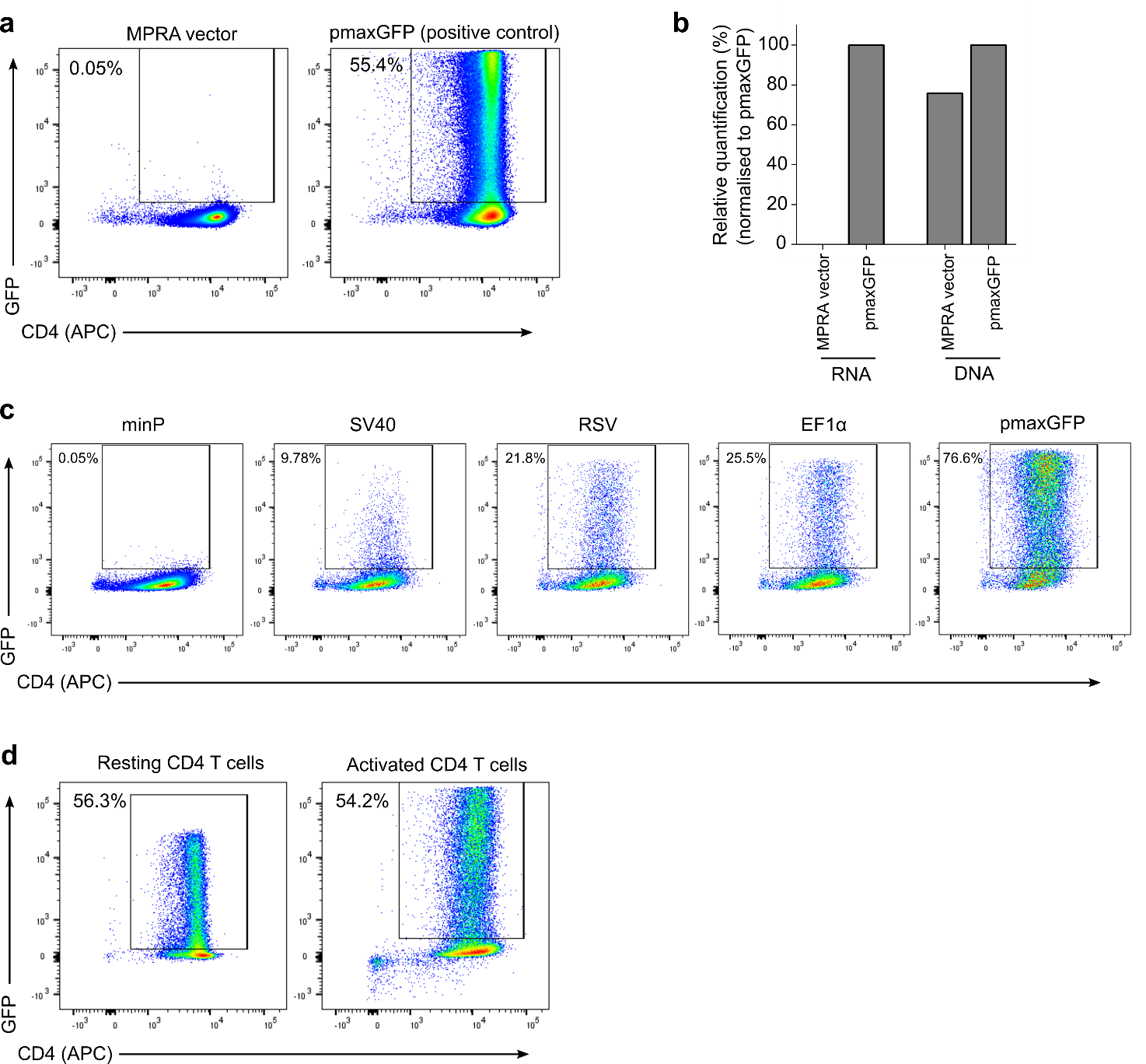

**Figure S1. Minimal promoter-based MPRA does not work in primary CD4 T cells, but can be successfully adapted using a stronger promoter.**

**a** Nucleofection of a minimal promoter-based MPRA vector into primary CD4 T cells does not lead to detectable GFP expression after 24 hours, unlike the positive control vector (pmaxGFP). **b** 24 hours after transfecting a minimal promoter-based MPRA vector into primary CD4 T cells, the vector can be recovered from the cells – confirming successful transfection – but no GFP RNA is detectable. Quantification by qPCR. **c** Flow cytometric assessment of the activity of a series of alternate promoters in primary CD4 T cells – all assayed 24 hours after transfection of 2μg vector into 5M CD4 T cells. **d** Representative plots of GFP expression 24 hours after nucleofecting an adapted MPRA vector (containing the RSV promoter, 5μg) into resting and stimulated primary CD4 T cells.

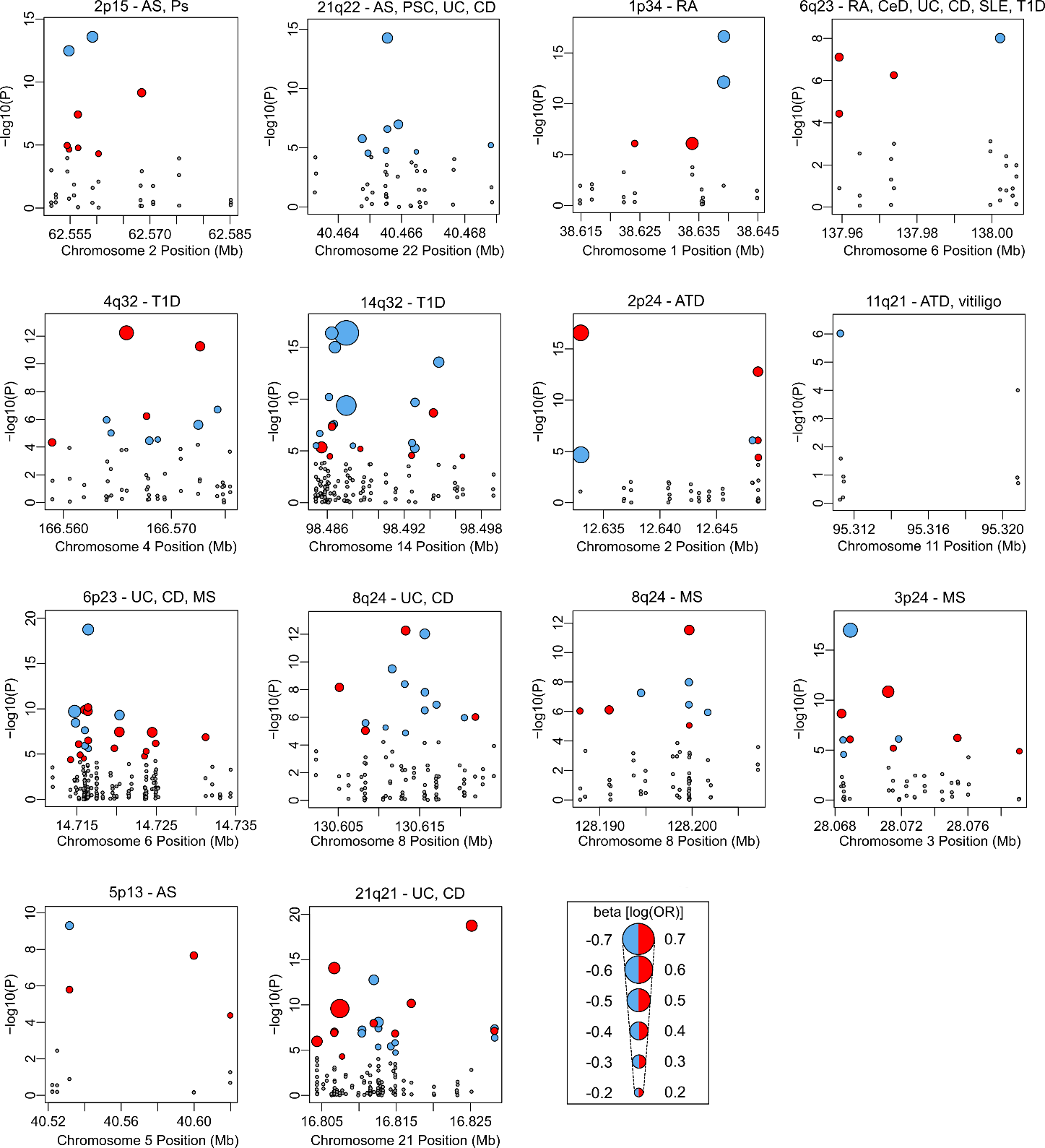

**Figure S2. Scaled Manhattan plots for 14 autoimmune disease associated loci – based on MPRA data from resting CD4 T cells.**

Scaled Manhattan plots of candidate SNPs in 14 autoimmune disease associated loci – based on expression-modulating effects in resting CD4 T cells. For SNP constructs with significant expression modulating effects (meta analysis P < 0.05/970) the size of each point is scaled to the effect size observed in the assay. The colour indicates the direction of the expression-modulating effect with respect to the risk allele. SNP constructs that did not pass this significance threshold are shown in grey. AS, Ankylosing Spondylitis; Ps, Psoriasis; PSC, Primary Sclerosing Cholangitis; UC, ulcerative colitis; CD, Crohn’s disease; RA, rheumatoid arthritis; CeD, coeliac disease; SLE, Systemic Lupus Erythematosus; T1D, Type 1 Diabetes; ATD, autoimmune thyroid disease; MS, multiple sclerosis.

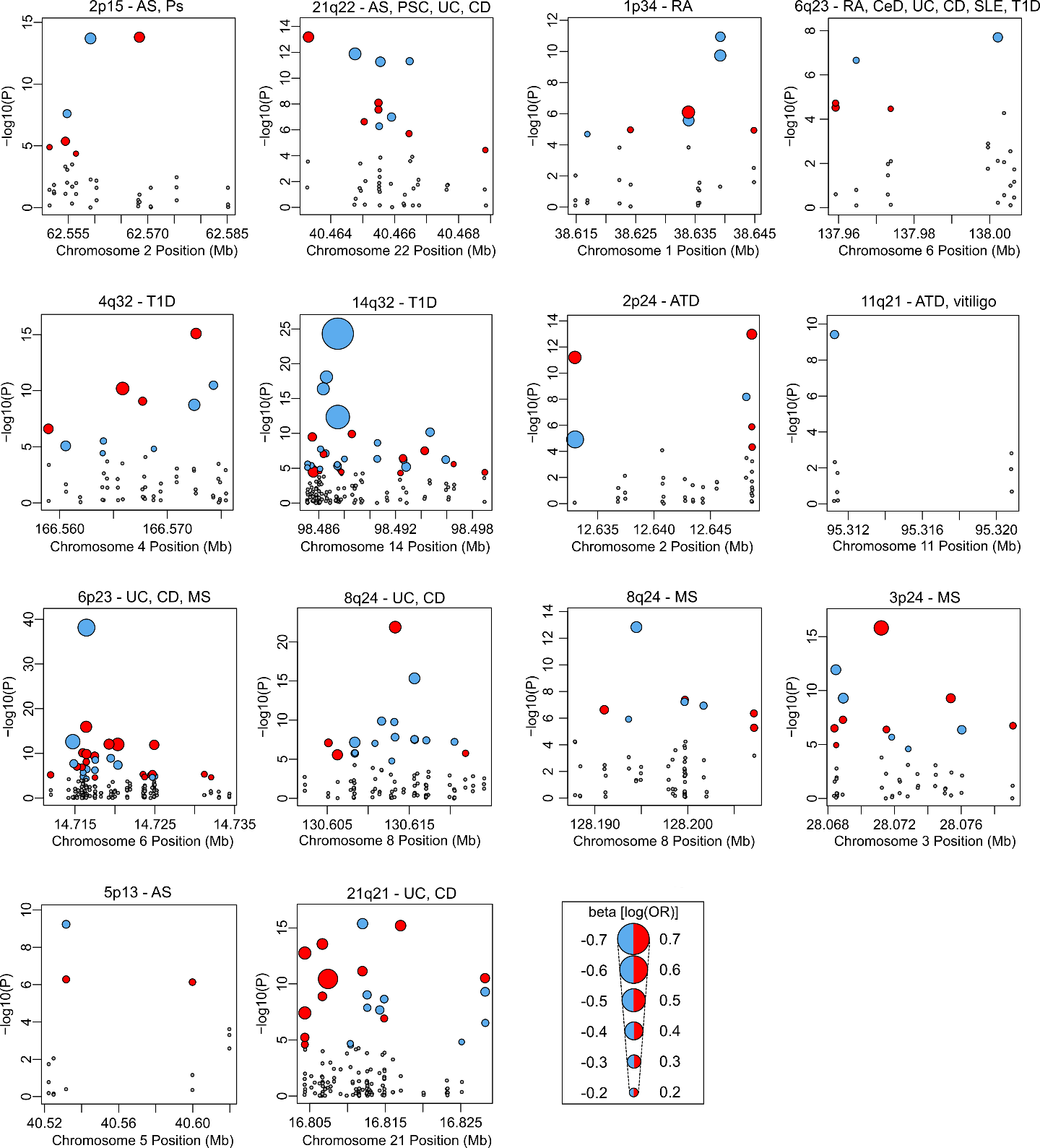

**Figure S3. Scaled Manhattan plots for 14 autoimmune disease associated loci – based on MPRA data from stimulated CD4 T cells.**

Scaled Manhattan plots of candidate SNPs in 14 autoimmune disease associated loci – based on expression-modulating effects in stimulated CD4 T cells. For SNP constructs with significant expression modulating effects (meta analysis P < 0.05/970) the size of each point is scaled to the effect size observed in the assay. The colour indicates the direction of the expression-modulating effect with respect to the risk allele. SNP constructs that did not pass this significance threshold are shown in grey. AS, Ankylosing Spondylitis; Ps, Psoriasis; PSC, Primary Sclerosing Cholangitis; UC, ulcerative colitis; CD, Crohn’s disease; RA, rheumatoid arthritis; CeD, coeliac disease; SLE, Systemic Lupus Erythematosus; T1D, Type 1 Diabetes; ATD, autoimmune thyroid disease; MS, multiple sclerosis.

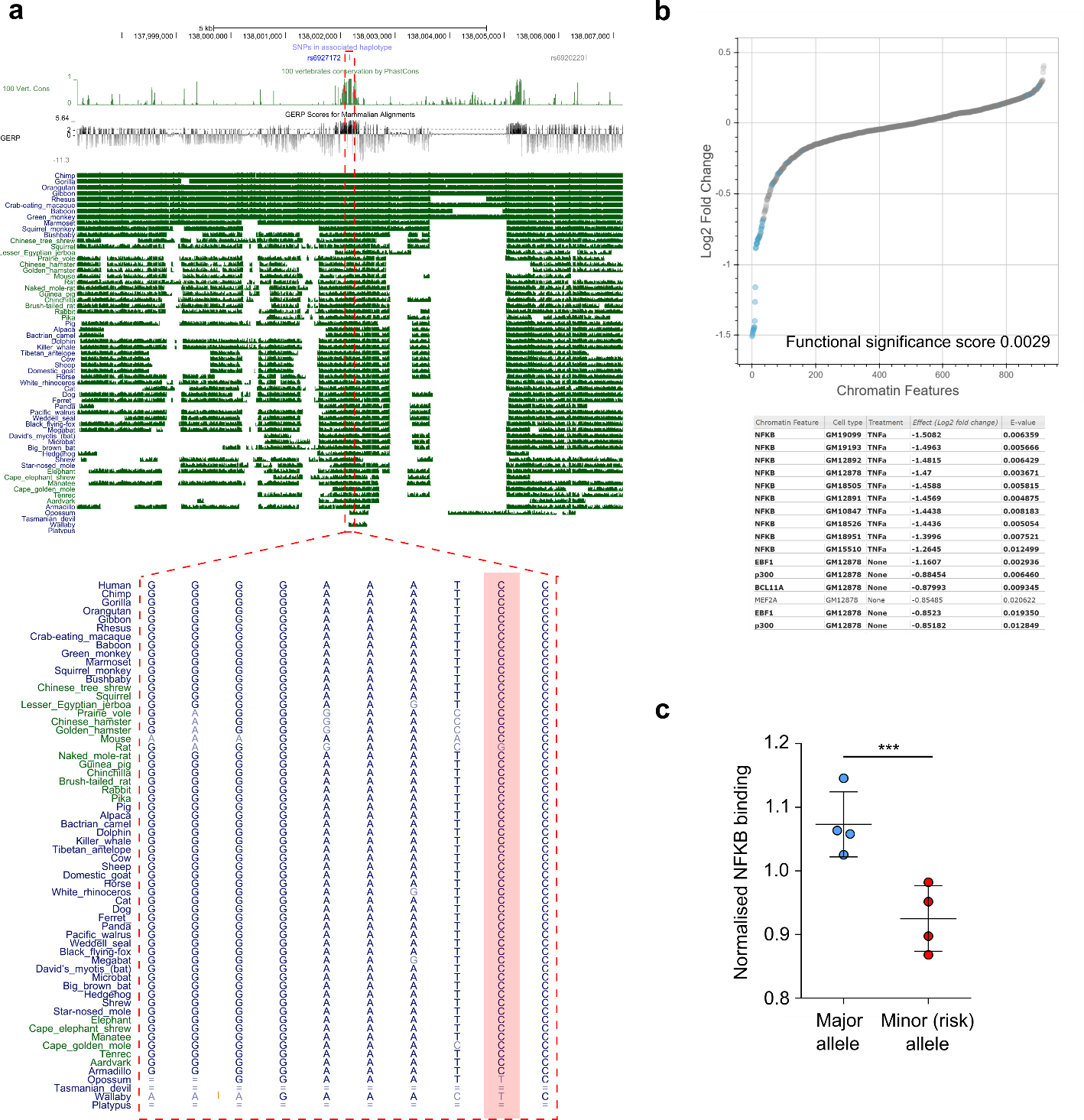

**Figure S4. rs6927172 lies in a highly conserved locus and is predicted to disrupt NF-κB binding.**

**a** Three analyses of conservation: PhastCons, Genomic Evolutionary Rate Profiling (GERP) and Multiz all show that rs6927172 lies in a highly conserved region. Data from UCSC Genome Browser. rs6927172 highlighted in pink in lower panel. **b** DeepSea analysis of candidate SNPs at 6q23 locus (using machine learning of regulatory sequence code from ENCODE chromatin-profiling data) predicts that rs6927172 is functionally significant with a significant effect on NF-κB binding. Inset table shows top results for rs6927172 ordered by effect size. **c** Following nucleofection of the MPRA vector library into primary CD4 T cells from 4 healthy individuals, cells were cross-linked and NF-κB immunoprecipitation was performed. Isolated plasmids were sequenced to assess for differential NF-κB binding. A SNP construct for rs6927172 showed significant allele-specific NF-κB-binding, with reduced binding to the risk allele containing vector.

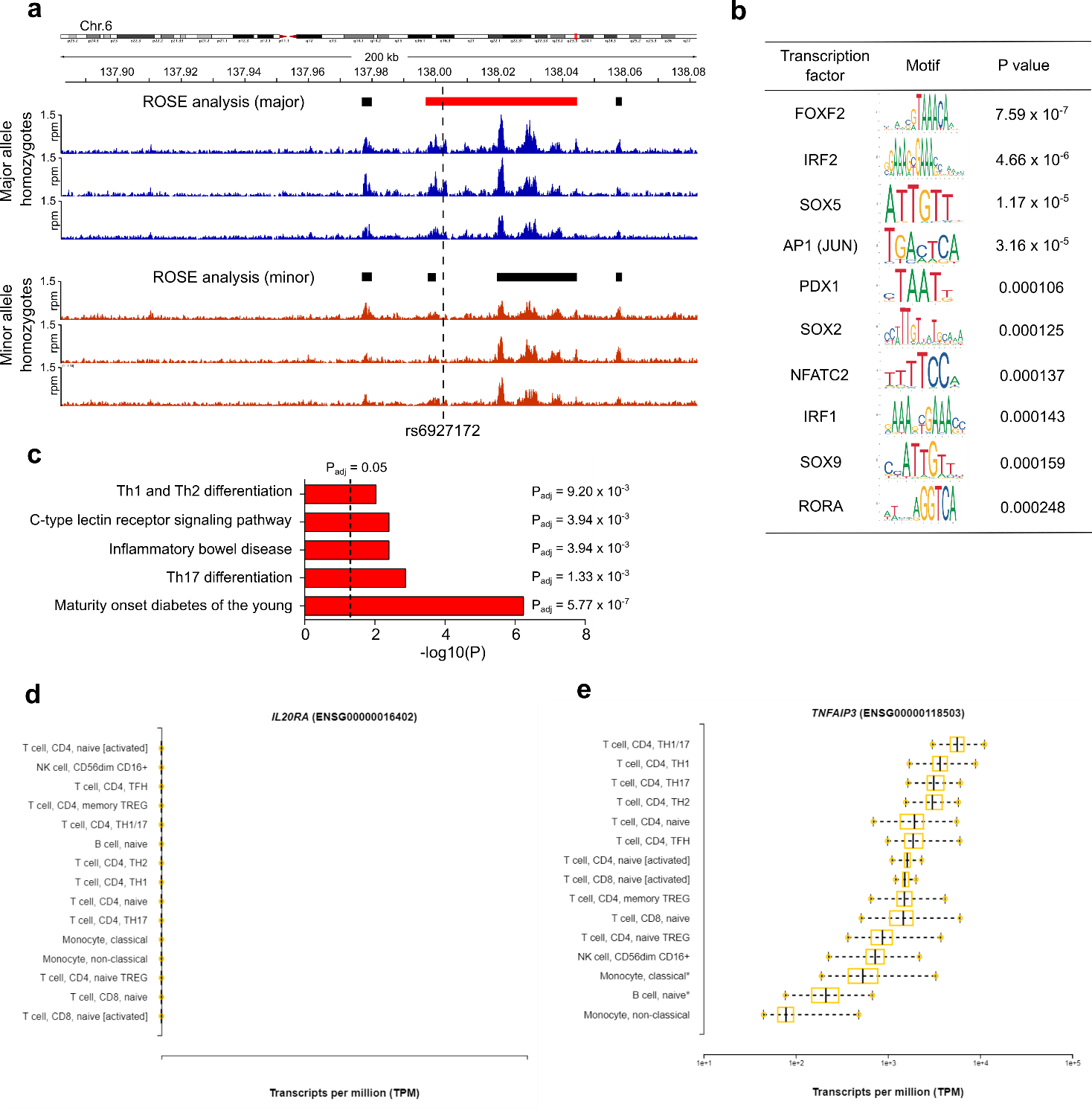

**Figure S5. rs6927172 disrupts a *TNFAIP3* super-enhancer at 6q23.**

**a** Normalised H3K27ac ChIP seq reads at the 6q23 locus in individual minor and major allele homozygotes at rs6927172. Bars indicate results from ROSE analysis (Ranking of Super-Enhancers, black = enhancer activity; red = super-enhancer). **b** The top 10 transcription factors whose binding motifs are over-represented within the constituent enhancer elements within the super-enhancer (compared to a background model based on all human promoters). Analysis performed using TRAP (TRanscription factor Affinity Prediction using Jaspar vertebrate matrices and Benjamini Hochberg correction for multiple testing). **c** Pathway analysis results (KEGG, FDR P < 0.01) using all 50 significantly-overrepresented transcription factors from TRAP analysis. Analysis performed using gProfiler. **d** **and** **e** Expression of *IL20RA* (**d**) and *TNFAIP3* (**e**) in primary immune cells from DICE database.

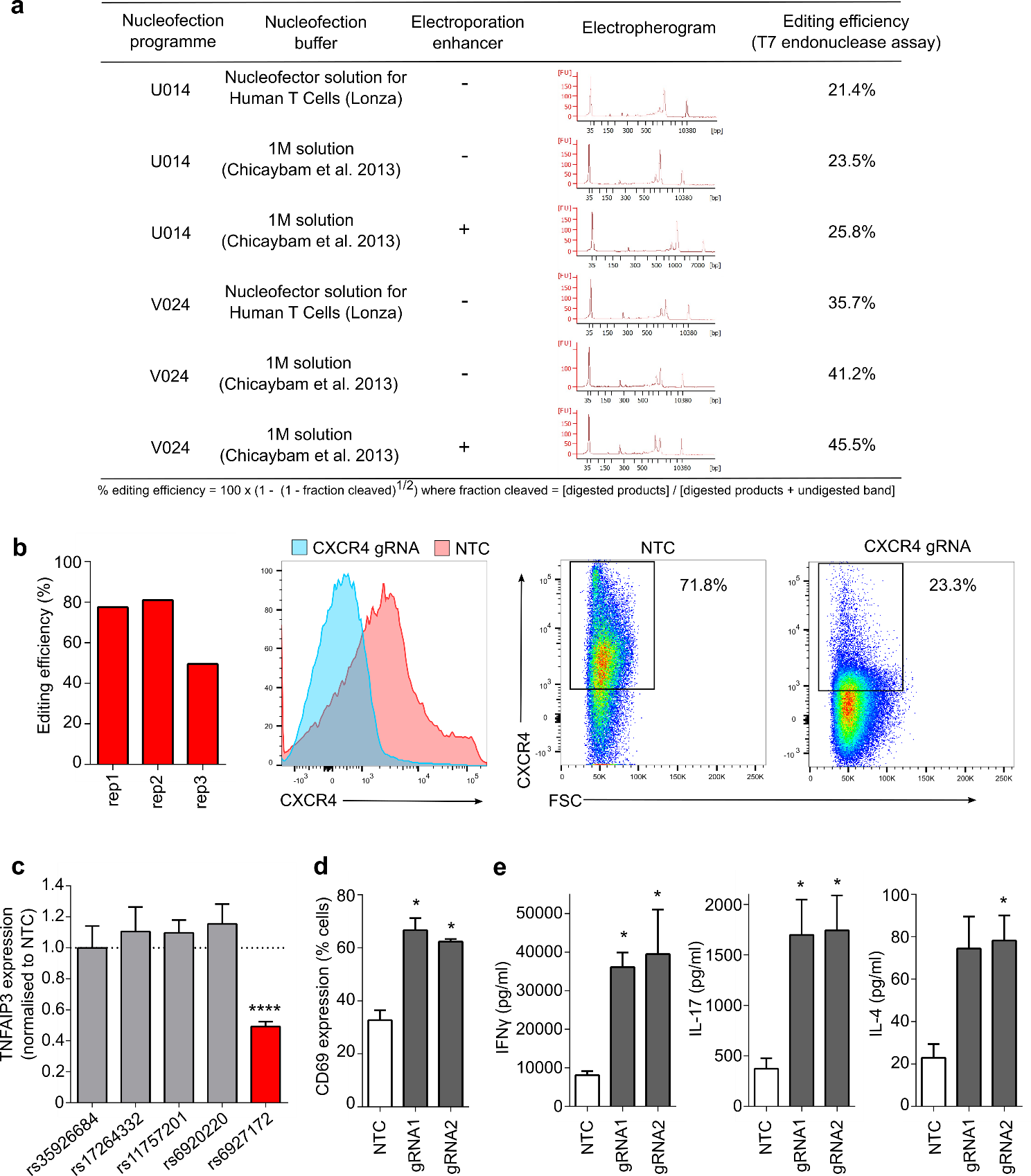

**Figure S6. CRISPR-Cas9 editing in resting primary CD4 T cells.**

**a** Optimisation of CRISPR editing in resting CD4 T cells using a Cas9 RNP containing positive control gRNA (targeting *HPRT*). On-target editing was assessed using a T7 Endonuclease assay. **b** Editing efficiency at the *CXCR4* locus assessed using ICE (left panel). Representative histograms and flow cytometry plots of CXCR4 expression on CD4 T cells following CRISPR editing (right panels). **c** *TNFAIP3* expression in EU-containing mRNA (EU added at time of stimulation) following individual deletions of candidate SNPs within the *TNFAIP3* super-enhancer (data from a minimum of 4 biological replicates). Mean indel rates: rs35926684, 55.0%; rs17264332, 54.2%; rs11757201, 52.0%; rs6920220, 72.1%; rs6927172, 64.5%). **d** Percentage of CD4 T cells expressing CD69, an activation marker, following CRISPR editing of *TNFAIP3* (n = 4, paired t-test, one-tailed). **e** Secretion of IFNγ, IL-17A and IL-4 following CRISPR editing of *TNFAIP3* in activated CD4 T cells (n = 4, paired t-test, one-tailed). Data represent mean +/- SEM. * P < 0.05; **** P < 0.0001.
